# Delineation of two multi-invasion-induced rearrangement pathways that differently affect genome stability

**DOI:** 10.1101/2023.03.15.532751

**Authors:** Diedre Reitz, Yasmina Djeghmoum, Ruth A. Watson, Pallavi Rajput, Juan Lucas Argueso, Wolf-Dietrich Heyer, Aurèle Piazza

## Abstract

Punctuated bursts of structural genomic variations (SVs) have been described in various organisms, but their etiology remains incompletely understood. Homologous recombination (HR) is a template-guided mechanism of repair of DNA double-strand breaks and stalled or collapsed replication forks. We recently identified a DNA break amplification and genome rearrangement pathway originating from the endonucleolytic processing of a multi-invasion (MI) DNA joint molecule formed during HR. Genome-wide sequencing approaches confirmed that multi-invasion-induced rearrangement (MIR) frequently leads to several repeat-mediated SVs and aneuploidies. Using molecular and genetic analysis, and a novel, highly sensitive proximity ligation-based assay for chromosomal rearrangement quantification, we further delineate two MIR sub-pathways. MIR1 is a universal pathway occurring in any sequence context, which generates secondary breaks and frequently leads to additional SVs. MIR2 occurs only if recombining donors exhibit substantial homology, and results in sequence insertion without additional break or SV. The most detrimental MIR1 pathway occurs late on a subset of persisting DNA joint molecules in a PCNA/Polδ-independent manner, unlike recombinational DNA synthesis. This work provides a refined mechanistic understanding of these HR-based SV formation pathways and shows that complex repeat-mediated SVs can occur without displacement DNA synthesis. Sequence signatures for inferring MIR1 from long-read data are proposed.

## Introduction

Structural variants (SVs) of the genome are the predominant class of driver mutations in most cancer types, and more numerous than the more widely studied single-nucleotide variants and indels (Cosenza et al. 2022; ICGC/TCGA Pan-Cancer Analysis of Whole Genomes Consortium 2020). These SVs fuel cancer onset and its evolution, promoting metastasis and chemoresistance (Notta et al. 2016). Abruptly acquired clusters of SVs (*i.e.* chromothripsis, chromoanagenesis and chromoplexy) were found in >20% of 2,658 patients in a pan-cancer analysis, and often play a causal role in cancer development (ICGC/TCGA Pan-Cancer Analysis of Whole Genomes Consortium 2020; Notta et al. 2016). The molecular process(es) underlying these complex SVs are incompletely characterized (Cosenza et al. 2022).

The maintenance of genome stability involves independent molecular mechanisms that inhibit the occurrence of DNA damage and achieve their accurate repair (Putnam and Kolodner 2017). Homologous recombination (HR) is a universal DNA damage repair and tolerance mechanism that uses an intact homologous dsDNA as a template. As such, it is often considered a high-fidelity repair mechanism. Yet, extensive experimental evidences have implicated HR in the formation of repeat-mediated SVs, an outcome exacerbated in various mutant contexts (Putnam and Kolodner 2017; Savocco and Piazza 2021). These SVs originate either (i) from unrestricted DNA synthesis initiated at an ectopic (*i.e.* non-allelic) displacement loop (D-loop) DNA joint molecule (JM) in a process called break-induced replication (BIR), or (ii) by endonucleolytic processing of ectopic JMs generated throughout the pathway (Savocco and Piazza 2021).

The tenet of BIR is a displacement DNA synthesis step that goes unabated until stabilization of the DSB end, either by telomere capture, merging with a converging replication fork, D-loop disruption and annealing of the extended ssDNA to a complementary DNA end, or by end-joining (Anand et al. 2013). BIR depends on processive displacement DNA synthesis by PCNA-Polδ in the context of a D-loop, which is reduced in the absence of Pol32 (Brocas et al. 2010; Donnianni and Symington 2013; Li et al. 2009; Lydeard et al. 2007; Saini et al. 2013; Donnianni et al. 2019; Payen et al. 2008). Consequences of BIR range from loss of heterozygosity to non-reciprocal translocation associated with a copy number gain, upon allelic and ectopic donor usage, respectively. More complex rearrangements may result from frequent template-switches over the course of BIR (Anand et al. 2014; Stafa et al. 2014). BIR is thus a HR-based mechanism that can generate complex, neo-synthesized chromosomal rearrangements from a single DSB end.

The role of structure selective endonucleases (SSEs) in the generation of HR-mediated chromosomal rearrangements has long been recognized as part of the canonical double-strand break repair (DSBR) model. Endonucleolytic resolution of an allelic inter-homolog double Holliday junction (dHJ) intermediate can lead to the formation of either a crossover or a non-crossover with equal likelihood. Resolution into a crossover can cause loss of heterozygosity of the centromere-devoid chromosomal region at the next cell generation (Szostak et al. 1983). If DNA strand invasion occurs at an ectopic repeat, a crossover can lead to various types of repeat-mediated SVs, including insertions, deletions, inversions, and translocations (Argueso et al. 2008; Sampaio et al. 2020). This vulnerability is not restricted to the resolution of dHJs, as other upstream JMs such as D-loops can also be processed by SSEs (Deem et al. 2008; Pardo and Aguilera 2012; Ruiz et al. 2009; Smith et al. 2009; Mazón and Symington 2013; Mazón et al. 2012).

We recently identified a genomic instability mechanism originating from “multi-invasions” (MI), a JM in which a single DSB end has invaded two independent dsDNA donors (Piazza et al. 2017) (**Fig. 1A**). MIs are readily formed in reconstituted *in vitro* reactions with ssDNA substrates of physiological length (*i.e.* >100 nt) and yeast and human Rad51, Rad54, and RPA proteins, and in *S. cerevisiae* cells (Piazza et al. 2017; Wright and Heyer 2014; Piazza et al. 2021b). They are presumably byproducts of homology search by inter-segmental contact sampling demonstrated for RecA (Forget and Kowalczykowski 2012). This tripartite recombination mechanism, termed multi-invasion-induced rearrangement (MIR), generates an oriented translocation between the two engaged donors in a HR- and SSE-dependent fashion (Piazza et al. 2017). MIR inserts the intervening sequence of the invading molecule between the two translocated donors. It occurs at varying frequencies depending on homology length and spatial location of the recombining partners, reaching frequencies of up to 1% in wild-type cells. Depending on the sequence context, it yields additional rearrangements between the DSB and donor molecules at high frequencies, from 15 to 85% of MIR recombinants (Piazza et al. 2017) (and see genome-wide approaches below). Finally, MIR relies only partially on Pol32, suggesting that at least a fraction of MIR events and/or the repair of newly generated breaks does not require extensive displacement DNA synthesis. Thus, MIR may generate punctuated bursts of SVs with no or limited DNA synthesis involved, unlike BIR.

**Figure 1:**
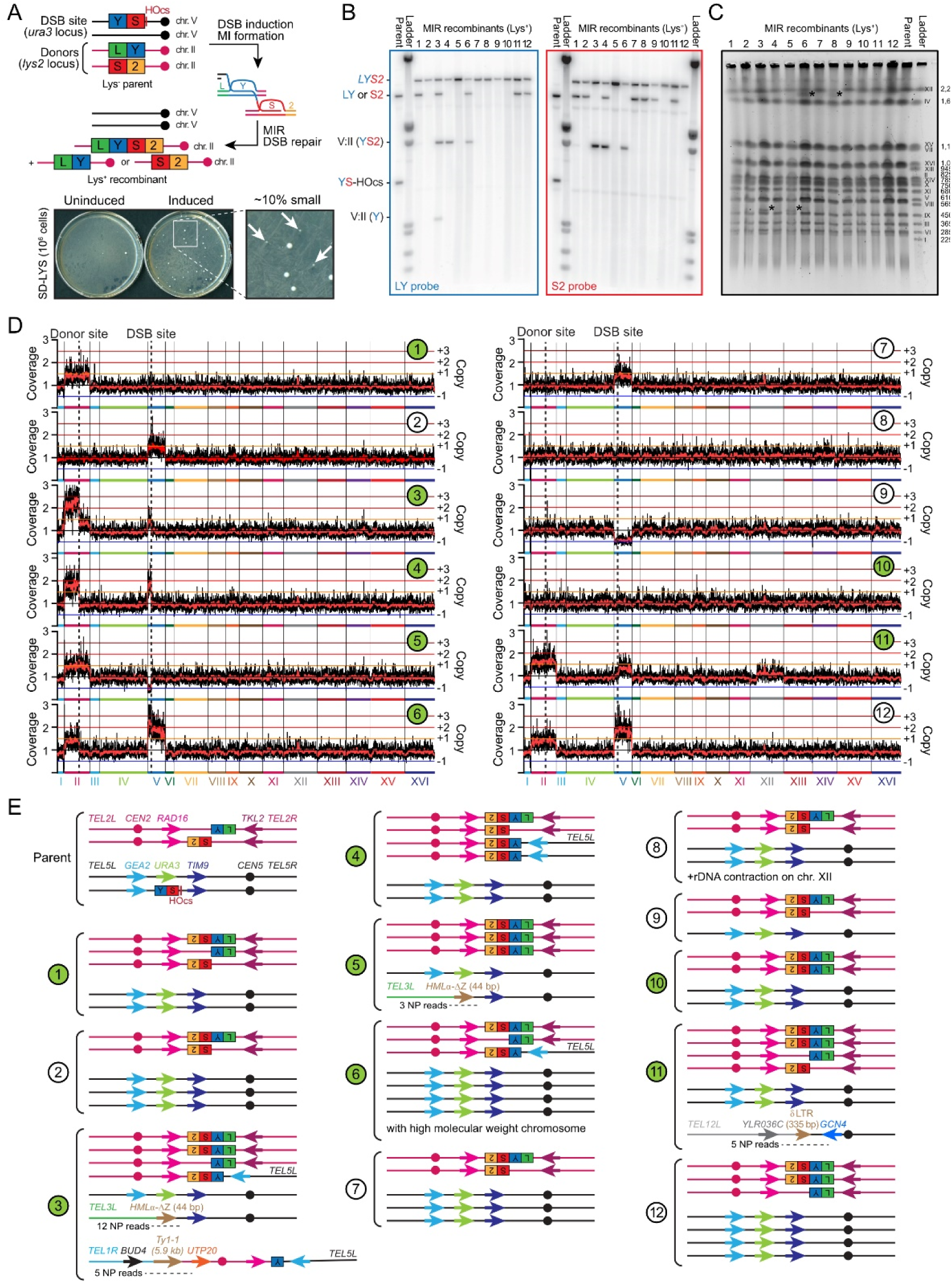
MIR frequently causes additional unselected rearrangements. (A) Experimental system in diploid *S. cerevisiae*. The heterozygous DSB-inducible *YS*-HOcs construct replaces *URA3* on chr. V. The *LY* and *S2* donors consist in the two halves of the *LYS2* gene (2090 and 2089 bp, respectively) at its locus on chr. II. This DSB-donor configuration is referred to as inter-chromosomal, with allelic donors. Translocation of the *LY* and *S2* donors restores a functional *LYS2* gene. Approximately 10% of induced Lys^+^ colonies are small on the initial plate, and exhibit a higher proportion of additional SVs than bigger colonies. Twelve small MIR recombinants were analyzed by Southern blot (B), PFGE (C), and high-throughput shotgun sequencing (D). Strains labeled in green were additionally analyzed by aCGH and Nanopore long-read sequencing. (C) The ladder corresponds to a *S. cerevisiae* strain from the YPH80 background, marginally different from our W303 parental strain. Ladder size is in kb. chr. V and chr. VIII co-migrate in the W303 background. *: chromosomal abnormality. (E) Deduced genome structure of the 12 MIR recombinants. The number of Nanopore (NP) reads encompassing the unselected rearrangements is indicated.

Here we investigated the existence of two MIR sub-pathways postulated on the basis of past molecular and genetic evidence: MIR1 that generates a translocation and additional DSB ends, applicable in any sequence context; and MIR2 that generates an insertion without forming an additional DSB but that has specific sequence requirements (see below, **Fig. 2A**) (Piazza and Heyer 2018; Piazza et al. 2017). We devised a highly sensitive molecular assay to quantify specific chromosomal rearrangements, which revealed that MIR1 occurs late on persistent DNA joint molecules enriched for MIs. Importantly, MIR1 occurs in the absence of Polδ-PCNA, unlike D-loop extension. These results elucidate a novel HR-based mechanism leading to tripartite SVs in the absence of displacement DNA synthesis.

**Figure 2:**
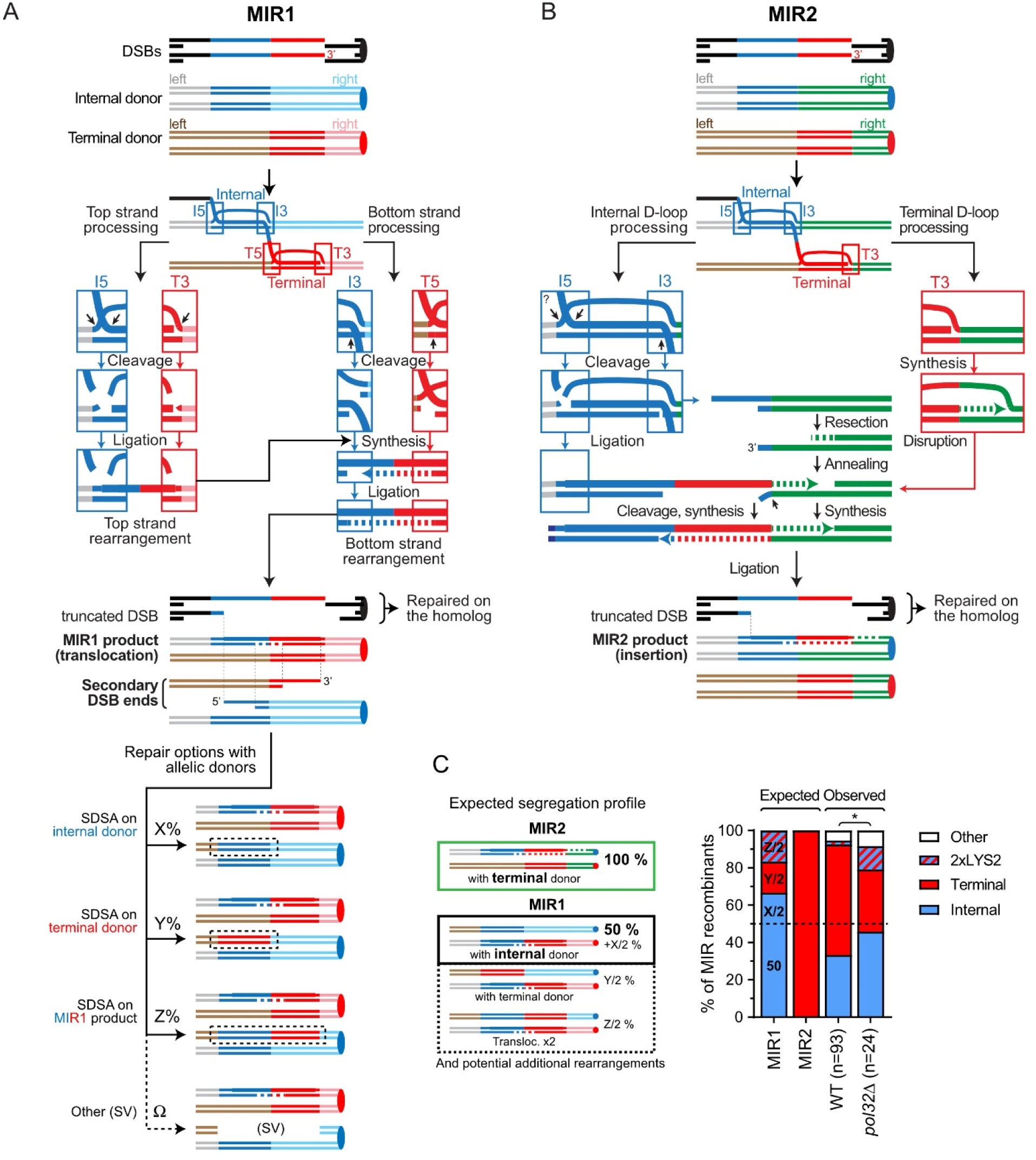
Models of multi-invasion-induced rearrangements. (A) MIR1 model. MIR1 generates a translocation and two additional single-ended DSBs. (B) MIR2 model. MIR2 generates an insertion. (C) Segregation products expected from MIR1 and MIR2, and of that observed in wild-type (APY89) and *pol32*Δ (WDHY4408) strains. Wild-type data are from Southern blot presented in (Piazza et al. 2017). Southern blot images for *pol32*Δ are shown in **Fig. S2C**. “Other” refers to strains bearing no donor and/or an SV involving one donor and the initiating DSB region. * denotes statistical significance (p <0.05, Fisher’s exact test).

## Results

### Experimental system

We previously developed an experimental system in diploid *S. cerevisiae* cells that specifically selects the MIR translocation product (Piazza et al. 2017, 2021b). This product, formed by a rearrangement that unites the two halves of the *LYS2* gene, leads to lysine prototrophy (Lys^+^ colonies). The system has three components: a site-specific DSB-inducible construct and two donors. The heterozygous DSB-inducible construct on chr. V consists of the HO cut-site (HOcs) and on one side the central part (***YS***) of the *LYS2* gene. This *YS* sequence has non-overlapping homology to two donors, the L**Y** and **S**2 halves of the *LYS2* gene (**Fig. 1A**). The Y and S share homologies are ∼1 kb-long each (**Fig. 1A**). In the reference strain used previously (Piazza et al. 2017), these donors were located at an allelic position on chr. II, in place of the original *LYS2* gene. In this heterozygous donor configuration, referred to as “allelic”, the sequences flanking the donors were identical. Other configurations were also used, in which the two donors were located at different, ectopic loci. These donor configurations were referred to as “ectopic”, with donors devoid of flanking homologies (see below). These latter configurations resulted in higher frequencies of secondary rearrangements (Piazza et al. 2017).

DSB formation at the HO cut-site is achieved with ∼99% efficiency within 1 hour of induction of *HO* gene expression (Piazza et al. 2017, 2021b). Its repair by gene conversion off the homolog takes up to 6 hours, during which MIR can occur (Piazza et al. 2021a, 2021b) (**Fig. 1A**). The basal and induced frequencies of Lys^+^ cells were determined for multiple independent cultures by plating cells on selective media prior to, and two hours post-DSB induction, respectively (**Fig. 1A**). In our reference wild-type strain, DSB induction causes a 100-fold increase in the frequency of Lys^+^ colonies, at 3.08 ± 0.3 x 10^−5^ colonies (Piazza et al. 2017) (see below).

### MIR frequently causes additional unselected rearrangements

Southern blot and quantitative PCR (qPCR) analysis of MIR products confirmed that all recombinants resulted in the formation of the *LYS2* gene at its expected locus, segregating with either the *LY* or *S2* donor. In a typical recombinant, the inducible DSB site is lost through gene conversion off the homologous chr. V (**Fig. 1A**) (Piazza et al. 2017). In ∼15% of total recombinants, additional, unselected rearrangements between the *Y-* and *S-*containing parts of our experimental system were observed in addition to the MIR product (Piazza et al. 2017). The prevalence of these additional rearrangements was much higher (∼90%) in small Lys^+^ colonies, which represent ∼10% of the recombinants (**Fig. 1A**). Here, we sought to determine MIR-associated SVs and copy number variation (CNV) genome-wide. To this end, 12 small Lys^+^ recombinants were analyzed by Southern blot (**Fig. 1B**), pulse-field gel electrophoresis (PFGE) (**Fig. 1C**) and paired-end short-read high throughput sequencing (**Fig. 1D**). A subset (1, 3, 4, 5, 6, 10, and 11) was further analyzed by array Comparative Genome Hybridization (aCGH) and long-read Nanopore sequencing. The structure of the recombinant chromosomes is depicted **Fig. 1E**.

The *LYS2* gene was restored at its locus in all (12/12) cases, as expected (**Fig. 1B**). This translocation segregated either with the *LY* or the *S2* donor in 8/12 cases (**Fig. 1B**). In 2/12 cases (#5 and 10) no remaining donor was visible, while in the last 2/12 cases (#1 and 11) the *LY* and *S2* donors were both present together with the *LYS2* translocation (**Fig. 1B**). Finally, 4 additional SVs involving chr. II and chr. V were visible in 3/12 recombinants (#3, 4, and 6). In total, 6/12 colonies (#1, 3, 4, 5, 6, 10) exhibited an additional chromosomal abnormality involving the DSB and/or the donors visible by Southern blot. Chromosome length determination by PFGE further revealed unambiguous chromosome abnormalities in 4/12 recombinants, in the form of either a ∼470 kb band (#3 and #5) or a ∼2 Mb band (#6 and 8) (**Fig. 1C**). Hence, 7/12 small Lys^+^ recombinants exhibited a detectable additional chr. II and/or chr. V abnormality together with the MIR translocation, consistent with past findings (Piazza et al. 2017).

High-throughput sequencing further revealed that all (12/12) of the analyzed small Lys^+^ MIR recombinants bore at least one additional chromosomal abnormality in addition to the selected MIR translocation. A total of 13 whole chromosome aneuploidies were detected in 9/12 strains analyzed, which all involved the chr. II and chr. V (**Fig. 1D**). The majority of additional SVs (5/9 instances in 3/4 strains) involved the *YS* break site on the chr. II and/or chr. V (**Fig. 1D**). However, in a substantial number of cases (4/9 instances in 3/4 strains) other chromosomes were also found to be involved, together with chr. II and/or chr. V. In colony #11, a translocation involving the right arm of chr. XII and the left arm of chr. V led to a 1,296 kb-long chromosome. The SV occurred between two ∼335 bp Solo δ elements sharing 89% homology: *YELWdelta6* from chr. V and *YLRCdelta3* from chr. XII (**Fig. 1E**). Colonies #3 and 5 exhibited a translocation between the right side of the DSB site on chr. V and the left side of *HML*α on chr. III, yielding a 474 kb chromosome visible by PFGE (**Fig. 1C**). The SV occurred at the 44 bp homology between the right side of the HO cut site (**Fig. 1E**). Finally, colony #3 exhibited an additional translocation between the right arm of chr. I and the left arm of chr. II. This latter translocation occurred at the position of two ∼5.9 kb Ty1-1 retrotransposons sharing 92% homology, *YARCTy1-1* on chr. I and *YBLWTy1-1* on chr. II. The resulting I:II translocated chromosome is expected to be 655 kb-long, but it was not observed by PFGE (**Fig. 1C**). Consequently, we favor the alternative possibility that the translocation occurred on the chr. II arm already involved in a chr. II:V translocation, yielding a I:II:V translocated chromosome of 433 kb migrating roughly at the same position as chr. IX (**Fig. 1E**). In all cases, substantial homologies (44 bp, 335 bp, and 5.9 kb) were present at these SV junctions. Finally, a distinct ∼2 Mb chromosome is visible by PFGE in recombinants #6 and 8 (**Fig. 1C**). Recombinant #6 bears the highest copy number gain of chr. V of all recombinants (**Fig. 1D**), which may give rise to this variant chromosome. Recombinant #8 does not exhibit chromosomal copy number change (**Fig. 1D**), but the ribosomal DNA (rDNA) coverage is reduced by ∼40% relative to the parental strain (**Fig. S1**). This band may thus correspond to a large heterozygous rDNA contraction on chr. XII.

In conclusion, detailed characterization of a subset of the ∼10% of small MIR recombinants confirmed that MIR frequently generates secondary rearrangements in addition to the primary selected translocation event (Piazza et al. 2017). It also revealed the involvement of other chromosomes together with those initiating MIR at sites of extensive homologies, suggesting a HR-dependent origin for these co-occurring rearrangements. The high frequency of whole chromosome aneuploidies also indicated that MIR is associated with chromosome mis-segregation. We sought to better characterize the MIR mechanism(s) leading to the formation of these secondary rearrangements.

### Existence of two MIR sub-pathways with different requirements for displacement DNA synthesis

We postulated the existence of two MIR pathways, MIR1 and MIR2, to account for (i) the segregation pattern of the *LY* and *S2* donors together with the MIR translocation and (ii) the presence of additional chromosomal rearrangements (**Fig. 2, Fig. S2A**) (Piazza and Heyer 2018). In the fully endonucleolytic MIR1 pathway, all four JM junctions are cleaved by SSEs (**Fig. 2A**). This pathway yields two additional one-ended DSBs, in addition to the translocation carried in place of the terminal *S2* donor (**Fig. 2A**). In an allelic donor configuration, repair of these secondary DSBs will yield an *LY*, an *S2*, or a second MIR product (*LYS2*), depending on the template used (**Fig. 2A**, detailed in **Fig. S2A**). This recombination product or the intact sister chromatid (internal *LY* donor) will segregate with the primary MIR translocation at the next cell division. If the donors are in an ectopic configuration, accurate repair is not possible and additional rearrangements are observed at high frequencies (see below and (Piazza et al. 2017)).

In contrast, MIR2 resolves the terminal invasion by synthesis-dependent strand annealing (SDSA), in which the extended end of the terminal D-loop pairs with the single-ended DSB generated upon cleavage of the internal D-loop (**Fig. 2B**). This pathway yields an insertion without additional DSBs. Consequently, the MIR2 product will segregate 100% of the time with the internal *LY* donor. No secondary rearrangement is expected.

The MIR1 and MIR2 mechanisms thus make clear predictions as to which donor segregates with the MIR translocation product (**Fig. 2C**). The observed segregation pattern, however, does not satisfy any single prediction (**Fig. 2C**), suggesting that both MIR1 and MIR2 mechanisms are at play with donors in an allelic configuration in wild-type cells (**Fig. 1A**). Although the MIR1 and MIR2 pathways have largely overlapping requirements (HR and SSE), they are expected to differ in:

- The processing of the terminal JM in the MI differs: it is cleaved in MIR1 and extended in MIR2. Moreover, the top strand of a MIR1 translocation can theoretically be produced without DNA synthesis (**Fig. 2A**).
- Their reliance on displacement DNA synthesis and 3’ homologies between the two donors differs, as displacement DNA synthesis is required only for MIR2 (**Fig. 2B**).
- A feature specific to MIR1 is that it generates two additional DSBs, which have the potential to lead to secondary rearrangements.

In the following sections we address the existence of the MIR1 and MIR2 pathways by challenging these predictions, and better define their relative contributions to the formation of chromosomal rearrangements in different sequence contexts.

### The absence of Pol32 causes a shift from the MIR2 to the MIR1 donor segregation profile

In order to gain insight into the mechanisms at play in the allelic donor situation and address the prediction that MIR2 requires displacement DNA synthesis, we analyzed MIR recombinants obtained in a *pol32*Δ mutant, defective for long-range recombination-associated DNA synthesis. *POL32* deletion caused a significant 2.5-fold decrease in MIR frequency (**Fig. S2B**) (Piazza et al. 2017). Southern blot analysis of 24 recombinants revealed a significant shift in the donor segregation pattern (**Fig. 2C, Fig. S2C**). Notably, the expected MIR2 signature (*LYS2*+*S2*) decreased from 55/93 to 8/24 (p = 0.037, Fisher’s exact test), with a compensatory increase in MIR1 signatures (*LYS2*+*LY* and 2x*LYS2*). The proportion of *LYS2*+*LY* recombinants reached 45.8%, approaching the minimal proportion expected for MIR1-only events (50%; **Fig. 2C**). The share of strains exhibiting additional chromosomal abnormalities also increased, as expected for a larger fraction of MIR1 events. These results are consistent with our inference that both MIR1 and MIR2 contribute to MIR translocations. They support our first prediction that MIR2 relies on displacement DNA synthesis. However, because significant displacement DNA synthesis can occur in the absence of *POL32* (McVey et al. 2016), it was unclear from this analysis whether MIR1 also requires displacement synthesis (see below).

### MIR2, but not MIR1, requires long homologies downstream of the donors

Our models posit that MIR2, but not MIR1, requires homology flanking the donors at their 3’ side. To address this specific prediction, we moved the terminal (*S2*) donor to an ectopic location on chr. V (*CAN1* locus, 83 kb away from the DSB site at *URA3*, see **Fig. 3A**) (Piazza et al. 2017). In this configuration referred to as “ectopic-trans”, the donors do not share extensive flanking homologies (only the 70 bp *LYS2* terminator), unlike in the initial allelic configuration. Only MIR1 is expected in this configuration, whose outcome is a chr. II:V translocation yielding an ∼895 kb chromosome with *LYS2* present in place of the *S2* donor at *CAN1*. We then restored substantial flanking homologies (500 and 1000 bp, from the initial 70 bp) at the 3’ side of the donors by modifying the *S2* donor (**Fig. 3A**). Restoration of this 3’ flanking homologous region is expected to enable MIR2, whose outcome is *LYS2* in place of the *LY* donor without associated translocation. Hence, MIR1 and MIR2 products can be straightforwardly distinguished.

**Figure 3:**
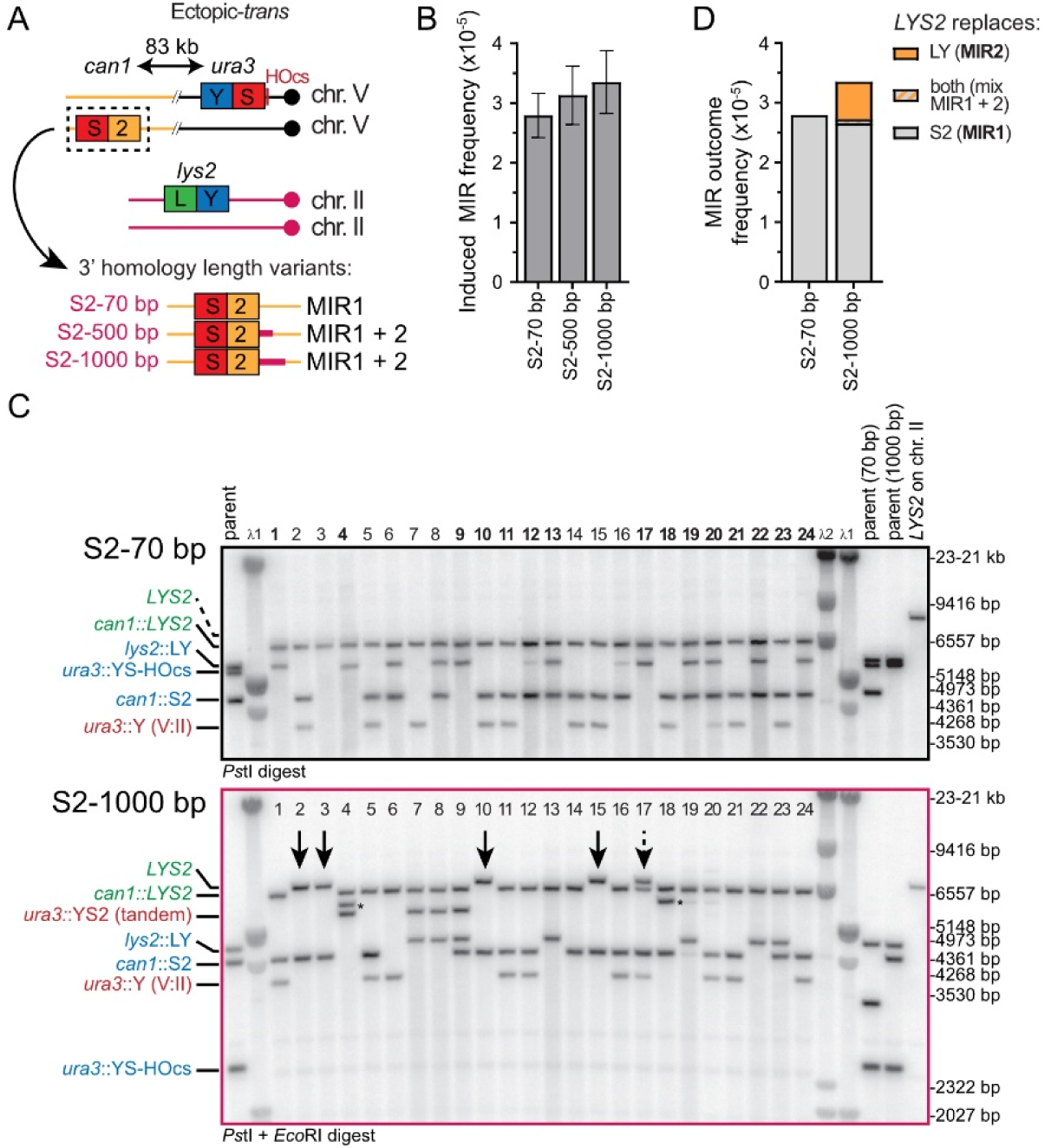
MIR2 uniquely requires homology at the 3’ side of each donor. (A) Experimental system to address the existence of MIR1 and MIR2 mechanisms. The *LY* and *S2* donors are ectopically located. The *S2* donor is in *trans* relative to the DSB site, on the chr. V homolog. Homology to the 3’ flanking sequence of the *LY* donor is increased near the *S2* donor from 70 bp to 1000 bp (purple region) corresponding to chr. II sequence. This extension does not contain full-length open-reading frames. (B) MIR frequency in wild-type cells bearing 70 bp, 500 bp and 1000 bp of homology 3’ of the LY and S2 donors (APY85, APY86, APY87, respectively). (C) Southern blot analysis of 24 Lys^+^ recombinants obtained with either 70 bp (n=46) or 1000 bp (n=48) of 3’ homology. For details on substrates and products length, see **Fig. S3A**. Remaining colonies are presented in **Fig. S3B**. Arrows indicate MIR2 products. The dashed arrow indicates a mixed colony. (D) MIR outcome frequencies.

Addition of flanking homology 3’ of the *S2* donor led to a modest, non-significant increase in MIR frequency (**Fig. 3B**). We determined the structure of the recombinants by Southern blot analysis on 46 and 48 Lys^+^ colonies obtained with the *S2*-70bp and the *S2*-1000bp donor constructs, respectively (**Fig. 3C** and **S3B**). Different restriction digestions were performed with each construct to be able to unambiguously distinguish parental molecules as well as primary and secondary rearrangement products (see **Fig. S3A** for a description of the various products). With the *S2*-70bp construct, all *LYS2* products were in the form of a chr. II:V translocation that replaced the *S2* donor at *CAN1*, as expected for a MIR1-only mechanism (**Fig. 3C** and **S2B**). PFGE analysis of a subset of 24 recombinants confirmed the presence of a ∼900 kb neo-chromosome in all cases (**Fig. S3C-D**), consistent with the length expected for such a chr. II:V translocation (see more below).

With the *S2*-1000bp construct, however, *LYS2* was found in place of the *LY* donor on chr. II in 9/48 cases, as expected for MIR2 products (arrows on **Fig. 3C** and **S2B**), significantly different from the 0/46 observed with the *S2*-70bp donor (p = 2.5×10^−3^, two-tailed Fisher’s exact test). 38/48 of the remaining colonies had *LYS2* in place of the *S2* donor, typical of MIR1, and 1/48 was a mixed colony (**Fig. 3C** and **S2B**). Thus, providing 3’ flanking homology between the donors is sufficient to restore MIR2. Combining these frequencies with total MIR frequency data indicates that these MIR2 events did not occur at the expense of MIR1 events, but were in addition to them, at a ∼5-fold lower frequency than MIR1 events (∼5.10^−6^ versus 2.7.10^−5^, respectively) (**Fig. 3D**). These observations corroborate our second prediction that MIR2, but not MIR1, requires substantial homology between the donors’ 3’ flanking sequences.

### MIR1, but not MIR2, is frequently associated with secondary rearrangements

MIR recombinants were subclassified based on their category (MIR1 or MIR2), the segregating donor(s) (A to D), and the secondary rearrangements detected (0 to 3), to yield classes in the form of, *e.g.* 1-C1 (**Fig. 4A**). First, MIR1 events obtained with *S2*-70bp and *S2*-1000bp did not exhibit significant difference in their class distribution (**Fig. 4B**). Half the recombinants exhibited at least one secondary chr. II:V translocation, and 39-41% exhibited a donor CNV (class A or D) (**Fig. 4B**). In total, 78.3% and 78.9% of MIR1 recombinants originating from *S2*-70bp or *S2*-1000bp-containing strains exhibited at least one additional chromosomal abnormality, respectively (Piazza et al. 2017). In contrast, only 1/9 MIR2 translocants (#33) contained an additional SV (**Fig. S3B**). This significant difference (p < 3.10^−4^ in both cases, two-tailed Fisher’s exact test) fulfills the last prediction that MIR1, but not MIR2, is associated with frequent secondary rearrangements. In conclusion, the three predictions of our two MIR models were satisfied:

- MIR2 specifically requires flanking homology on the 3’ side of the donors, but not MIR1 (**Fig. 3**).
- MIR2 requires sufficiently long displacement DNA synthesis tracts to make it partly dependent on Pol32 (**Fig. 2C**).
- Additional rearrangements occur mainly through MIR1, which differs from MIR2 in that it gives rise to two single-ended DSBs in addition to the MIR product (**Fig. 3**).

**Figure 4:**
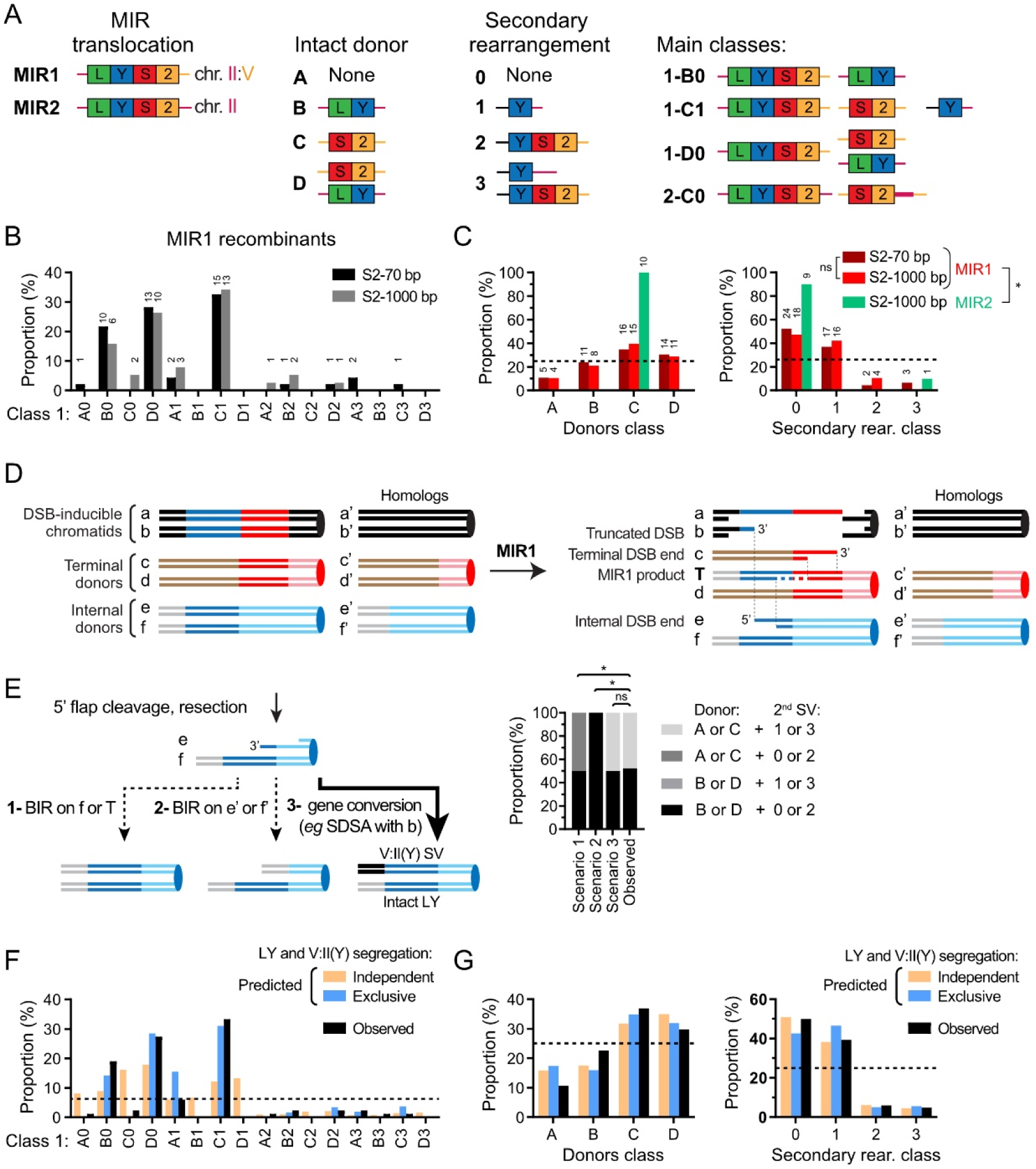
MIR1 is frequently associated with additional structural rearrangements, unlike MIR2. (A) Classification of Lys^+^ recombinants, based on MIR type, remaining donor(s), and additional rearrangement involving the donor and the break site. The most frequent MIR classes are depicted. (B) Distribution of MIR1 events obtained with the *S2*-70bp (n=46) and *S2*-1000bp (n=38) donor constructs. (C) Distribution of donor segregation (left) and secondary rearrangements (right) in MIR1 and MIR2 events obtained with the *S2*-70bp and *S2*-1000bp donor constructs. Number of events indicated above each bar in (B and C). (D) Expected location of the single-ended DSBs generated on the internal (*LY*) and terminal (*S2*) donors. Each chromatid is labeled. Note that in the ectopic-*trans* donor configuration, a/b are equivalent to c’/d’ and c/d are equivalent to a’/b’. They are not shown connected for the sake of simplicity and to avoid inducing uncertain associations between the repair of the initiating DSB ends and of the secondary DSB generated at the terminal donor. (E) Left: Three scenarios for the repair of the single-ended DSB at the internal (LY) donor. Right: the expected donor and SV classes for each scenario, and the observed classes distribution. * p<0.05, χ^2^ contingency test. (F) Predicted MIR1 class distribution assuming either independent or exclusive segregation of the *LY* donor and the V:II(*Y*) SV, and observed class distribution of the pooled *S2*-70bp and *S2*-1000bp-containing strains (n = 84). (G) Same as F, with distribution pooled by class (left) and secondary rearrangements (right).

### Analysis of secondary rearrangements in MIR1 recombinants

Finer MIR1 class analysis revealed specific associations or exclusions between additional DSB:donor SVs and donor retention (**Fig. 4B** and **Fig. S4**). Three main classes emerged: B0, C1 and D0 accounting for 19, 33 and 27% of total MIR1 recombinants, respectively (67/84 total, *S2*-70bp and *S2*-1000bp pooled; **Fig. 4A** and **B**). High-throughput sequencing or aCGH analysis of a subset of 4, 6, and 5 such recombinants obtained with the *S2*-70bp donor confirmed the expected chr. II and chr. V CNVs for each of these classes and revealed two instances of additional unselected CNVs (**Fig. S5A-C**, and see below). Notably, the class 1 SV (*i.e.* V:II(Y)) is the most common single SV observed (33/38), and is primarily associated with a C donor class (C1 = 27/33). Conversely, retention of the *LY* donor (*i.e.* B and D classes) was never associated with a V:II(Y) SV (0/19 and 0/25, respectively; **Fig. 4B** and **Fig. S4A-B**). This distribution significantly departs from that of an independent assortment of *LY* donor (B0) with the II:V(Y) SV (C1), in which 5/84 B1 events would be expected (p = 0.02, χ^2^-test), and indicates that the two are mutually exclusive. This observation allows mechanistic inferences as to the repair of the primary and secondary DSBs generated at donor sites following MIR1 (**Fig. 4D**).

Repair of the one-ended DSB generated in the *Y* region of the *LY* donor (e in **Fig. 4D**) upon MIR1 can occur by: (i) BIR using the intact sister chromatid (f” in **Fig. 4D**), (ii) BIR using the intact homolog (e’ and f’ in **Fig. 4D**) following additional resection and 3’ flap cleavage, or (iii) gene conversion with the left side of the initiating DSB (a or b, **Fig. 4D-E**). Repair by BIR predicts donor class distributions starkly different from those observed (**Fig. 4E**). However, gene conversion followed by random segregation of the V:II(Y) translocation and the intact *LY*-containing chromatid will result in 50% MIR1 recombinants bearing the *LY* donor (B+D class) and 50% lacking it (A+C class), not significantly different from the observed donor class distributions (52.2% and 47.8%, respectively; **Fig. 4E**). Consequently, repair of the one-ended DSB generated on the internal donor predominantly occurs by gene conversion together with the *Y*-containing ends on chr. V (**Fig. 4D-E**). In contrast, MIR2 recombinants exhibited the C class only, confirming that none of the terminal donor was damaged and lost in the MIR2 process.

The terminal (*S2*) donor is retained in the majority (56/84) of MIR1 recombinants (C and D classes, **Figs. 4B-C**). Because they are located at the same position at the *CAN1* locus on separate chromatids of the same homolog, the MIR1 translocation and the *S2* donor are not expected to co-segregate (**Fig. 4D**). Co-segregation thus implies that they are frequently disjoined, upon inter-homolog crossover or formation of an additional SV in the *CAN1-CEN5* interval during the repair of the initiating or secondary DSBs (**Fig 4D**). For instance the V:II(Y) SV, which takes place at the *URA3* locus in the *CAN1-CEN5* interval, frequently co-segregates with the *S2* donor (C1 versus A1 class, **Fig. 4B**). The resulting 83 kb insertion of the *CAN1-URA3* interval at the *LYS2* locus leads to a duplication detected in all (5/5) the C1 clones analyzed by aCGH (**Fig. S5C**). Consistently, careful PFGE analysis revealed the slight size difference between the resulting II:V:II chromosome (∼927 kb) and the II:V chromosome produced by MIR1 (∼895 kb), which is associated with a decreased intensity of the chr. II band (**Fig. S3C-D**). Thus, through the repair of the secondary DSB “e”, the MIR1 translocation is disjoined from *CEN5* and therefore from the *S2* donor. Our experimental setup does not allow us to quantify inter-homolog crossover products resulting from repair of the initiating DSB with the chr. V homolog as a template. However, these products were observed in half of all cases in a similar system with unrestricted homologies (Ho et al. 2010), likely explaining the proportion of D0 over B0 class (23/84 and 16/84, respectively).

Finally, V:II(YS2) SVs are infrequently observed (9/84; SV class 2 and 3, **Fig. 4C**). This SV can result from a half-crossover between the *YS*-containing initiating DSB (a or b) and the *S2* donor. The co-occurrence of V:II(YS2) and V:II(Y) SVs (class 3: 3/84) is not significantly lower from that expected for independent occurrence of class 1 and class 2 SVs (∼5/84). Consequently, the formation and the segregation of these two SVs occurs without detectable interference. This finding implies that cells undergoing MIR1 experience a failure to faithfully repair both initiating DSBs using the homolog as a template.

In conclusion, independent treatment of the repair of the initiating DSB and of the one-ended DSBs generated on the internal (*LY*) and terminal (*S2*) donors largely recapitulates the observed donor and SV class distributions (**Fig. 4F** and **G**). However, it fails to account for the absence or very low proportion of certain classes (A0, C0, B1, and D1), which can be explained by exclusive segregation of the LY donor with the V:II(Y) SV (**Fig. 4F**). With this assumption, only class A1 remained 2.6-fold lower than predicted. Our analysis thus broadly defines the mechanisms leading to the formation of secondary rearrangements initiated by a MIR1 event and can explain their segregation patterns.

### The Chromosomal Rearrangement-Capture (CR-C) assay detects MIR1 translocation with high sensitivity

To study the kinetics of MIR and the requirements for essential factors we developed Chromosomal Rearrangement-Capture (CR-C), a highly sensitive molecular assay for detection of rare chromosomal translocation in a cell population (rationale see **Fig. 5A**). CR-C relies on the proximity ligation of unique restriction sites covalently united upon translocation of the top DNA strand (**Fig. 1A**; detailed experimental procedure and normalization see **Methods**) coupled with the high dynamic range of a quantitative PCR readout. We used it to detect MIR1 products in the ectopic donor configuration upon formation of a homozygous DSB (**Fig. 5A**). This unrepairable system purposefully prevents growth resumption of cells repairing the break early, and enables quantification of DNA JM and MIR1 product in a fixed number of cells (Piazza et al. 2019). No MIR1 product is detected prior to DSB induction or in Lys^−^ colonies recovered after DSB induction (**Fig. 5B**). An induced Lys^+^ colony containing a single translocation in a diploid genome exhibits the expected CR-C value of 0.5±0.022 (**Fig. 5B**). The MIR1 product detection is Rad51-dependent, as is MIR (Piazza et al. 2017) (**Fig. 5C**). Serial dilution of DNA from a MIR1-containing strain (induced Lys^+^) in the DNA of a MIR1-devoid strain (induced Lys^−^) show that CR-C can reliably detect translocation frequencies down to about 5×10^−5^ (**Fig. 5D**). Accordingly, MIR (Lys^+^) frequency and MIR1 levels per haploid genome equivalent 24 hours post-DSB induction are in good quantitative agreement (**Fig. S6**). The CR-C assay is thus a sensitive molecular assay for the detection of rare translocations in a cell population.

**Figure 5:**
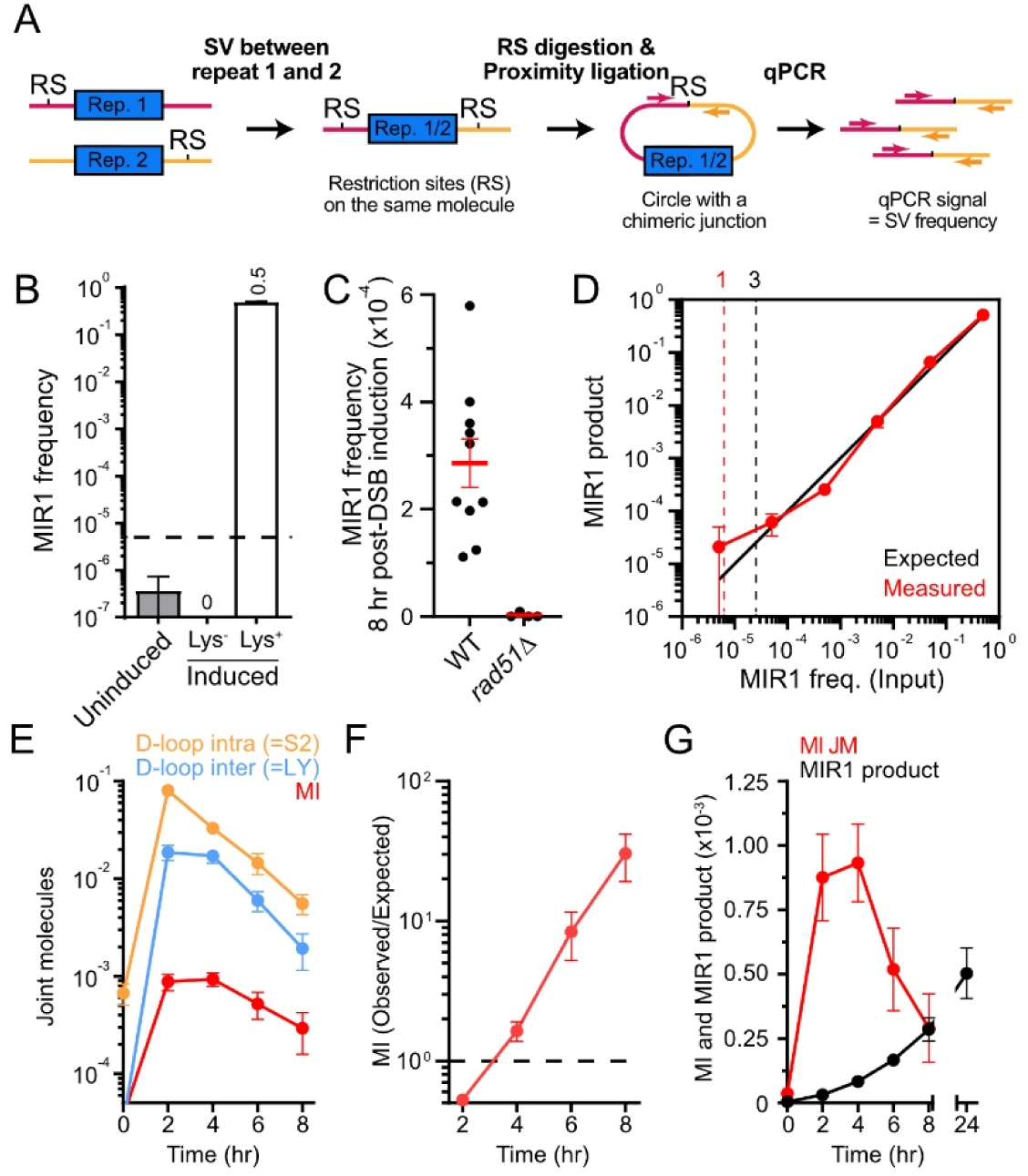
Chromosomal Rearrangement-Capture (CR-C) reveals the late occurrence of MIR1. (A) Rationale of the CR-C assay for detection of repeat-mediated SVs. (B) MIR1 signal in a wild-type strain (APY611) either uninduced, induced without having underwent MIR (Lys-), and in a selected Lys^+^ translocant containing a heterozygous MIR1 translocation. (C) MIR1 signal 8 hour post-DSB induction in a wild-type strain and in a *rad51*Δ mutant (APY625 and APY704, respectively). (D) Experimental determination of the sensitivity of the CR-C assay. DNA from a diploid strain containing a MIR1 translocation (Lys^+^) was serial diluted in the DNA of a Lys^−^ strain and the CR-C process performed. Dotted lines indicate the threshold for 1 and 3 molecules expected on average per qPCR reaction. (E) Quantification of D-loop JMs at the *LY* and *S2* donors and of MI following DSB formation in wild-type cells (APY625). (F) Ratio of the observed MI over those expected based on the product of independent LY and S2 D-loops. (G) Kinetics of MI abundance and MIR1 translocation following DSB formation.

### MIR1 occurs at late time points on a subset of persisting MI JMs

The D-loop-Capture (DLC) (Piazza et al. 2017, 2019, 2021b; Reitz et al. 2022) and CR-C assays enabled the determination of the temporal succession of D-loops, MIs, and MIR1 product formation following DSB induction (**Figs. 5E-G**). Improvements to the DLC protocol further enabled estimating the absolute D-loop and MI levels per haploid genome equivalent (**Methods**) (Reitz et al. 2022). D-loops at the intra-chromosomal S2 donor were highest at the earliest time point assayed (2 hours) and decreased monotonically 2- to 3-fold every 2 hours (**Fig. 5E**). D-loops at the inter-chromosomal LY donor peaked 4 hours post-DSB induction, before declining at 6 and 8 hours (**Fig. 5E**). These D-loop formation/decay kinetics were similar to those previously reported at these loci in haploid cells (Piazza et al. 2019, 2021a). The kinetics of MI formation followed that of the rate-limiting invasion of the inter-chromosomal donor (**Fig. 5E**). Average intra- and inter-chromosomal D-loops amounted to 3.3% and 1.7% of broken molecules 4 hours post-DSB induction. MI formed infrequently, involving 0.093% of broken molecules. This observed quantity of MIs closely matched that expected assuming independent invasion of the intra and inter donors 2 and 4 hours post-DSB induction, with an observed/expected ratio of 0.53 and 1.64, respectively (**Fig. 5F**). Though MIs declined in absolute terms over time, their observed/expected ratio gradually increased past the level expected from independent invasion events (**Fig. 5F**). Approximately 5.3% and 15.1% of intra and inter D-loops were thus part of a MI at 8 hours post-DSB induction, respectively. Hence, the subset of cells with JMs persisting past 4 hours post-DSB induction are enriched for MIs.

MIR1 occurred in a delayed fashion relative to both individual D-loops and MI formation (**Fig. 5G**). The increase in MIR1 product mirrored the decrease in MIs from 4 to 8 hours, with little MIR1 occurring afterwards (**Fig. 5G**). In absolute terms, the total MIR1 products observed at 8 hours (2.8×10^−4^) amounted to only a fraction of the MIs detected at any given time point (from 9.3×10^−4^ at 4 hours to 2.9×10^−4^ at 8 hours; **Fig. 5G**). Consequently, only a subset of MIs is converted into a MIR1 product, but this fraction increases over time. This result suggests that the pathway leading to a MIR1 product from a decreasing pool of MI intermediates mainly occurs at late time points relative to DSB induction. MIR1 resolution thus broadly coincides with the timing of DNA damage checkpoint adaptation (Toczyski et al. 1997).

### MIR1 does not require displacement DNA synthesis

The translocation of the top strand by MIR1 is predicted to be achieved in an endonucleolytic fashion (**Fig. 2A**). Restoration of the complementary strand will involve DNA synthesis using the ssDNA as a template. Accordingly, DNA synthesis that requires displacement of a complementary strand (displacement DNA synthesis) by Polδ-PCNA (Donnianni et al. 2019; Li et al. 2009) is predicted to be dispensable for MIR1 (**Fig. 2A**). To address this prediction, we depleted the Polδ catalytic subunit Pol3 alone or in combination with the PCNA clamp loader subunit Rfc1, by coupling an auxin-induced protein degradation and a Tet-Off transcriptional repression system (Donnianni et al. 2019; Kulkarni et al. 2020; Tanaka et al. 2015). As expected, depletion of Pol3 is lethal (**Fig. S7A**). Addition of doxycycline and auxin resulted in rapid elimination of both proteins (**Fig. S7B-C**), allowing analysis of their function specifically during DSB repair. Measurement of D-loop extension over 400 bp using our D-loop extension (DLE) assay (Piazza et al. 2018) revealed that the tagged Pol3 and Rfc1 proteins did not affect recombination-associated DNA synthesis. However, co-depletion of Pol3 and Rfc1 caused a >10-fold reduction in recombination-associated displacement DNA synthesis (**Fig. 6A**), an expected yet never demonstrated outcome. Importantly, Pol3/Rfc1 depletion did not significantly affect MIR1 product formation (**Fig. 6A**). Presence of the tagged proteins caused a ∼2-fold increase in MIR1. Joint molecules quantification by DLC revealed that presence of either or both tags, and to a lower extent protein depletion, caused an increase in the amounts of individual D-loops and MIs (**Fig. 6A** and **Fig. S7D**). This increase presumably results from a defect in Rfc1-dependent PCNA recruitment, which fails to direct D-loop disruption by Srs2 (Piazza et al. 2019; Robert et al. 2006; Miura et al. 2013; Liu et al. 2017; Burkovics et al. 2013). Consequently, we repeated these experiments in a *RFC1^+^*strain. Depletion of Pol3 alone reduced D-loop extension by ∼3-fold, without substantial change to D-loops and MI levels compared to a wild-type strain (**Fig. 6B** and **Fig. S7D**). It suggests that Rfc1 tagging and depletion, but not Pol3, caused elevated JM levels in the *rfc1-AID pol3-iAID* strain. Importantly, Pol3 depletion did not affect MIR1 (**Fig. 6B**). These results fulfill the final prediction that MIR1 occurs in the absence of displacement DNA synthesis.

**Figure 6:**
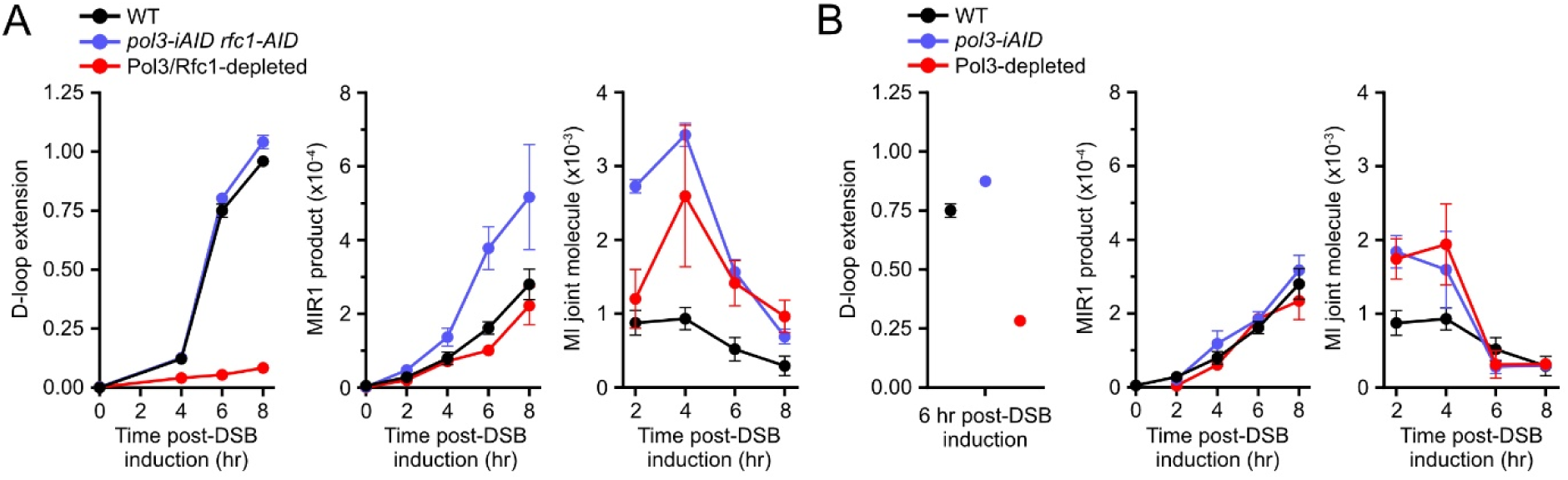
MIR1 occurs in the absence of Pol3 and Rfc1. (A) D-loop extension (left), MIR1 product formation (center), and MI joint molecules (right) in a *pol3-iAID rfc1-9Myc-AID* strain (WDH6066) strain upon mock-treatment or inhibitor treatment. (B) Same as in (A) in a *pol3-iAID* strain (WDHY6065). Data are mean±SEM. All data are n≥3 biological replicates, except for *pol3-iAID* (n=2 for CR-C and DLC; n=1 for DLE).

## Discussion

### Delineation of two MIR sub-pathways and their specific requirements

Here we provide genetic and molecular evidence for two MIR sub-pathways, their respective sequence and protein requirements, and their likelihood of generating SVs. MIR1 utilizes endonucleolytic resolution, without a need for shared sequence homology between the donors (**Fig. 2A**). As such, it can occur between any two genomic sites and thus represents a potentially ubiquitous repeat-mediated SV formation mechanism. MIR1 results in two additional DSBs, which can be accurately repaired only if there is homology between the two donors or their surrounding regions, as is the case at allelic sites (**Fig. S2A**). In the absence of such homology, there is no opportunity for accurate repair of these ends, and SVs frequently form involving the initiating and the donor sites (**Fig. 4**), as well as other sites exhibiting substantial levels of homology (**Fig. 1**). We could deduce the pathway leading to frequent secondary SVs, such as a chr. V:II SV, produced by gene conversion or SSA between the cleavage product of the internal invasion and one of the initiating DSB ends (**Fig. 4E**). Finally, MIR1 occurs in the absence of Polδ-PCNA, indicating that the large-scale rearrangements associated with this pathway involving multiple DNA repeats can happen without displacement DNA synthesis, in a endonucleolytic fashion. This MIR1 pathway contrasts with SV formation by repeat-mediated template-switches during BIR and fork restart, which are dependent on extensive displacement DNA synthesis (Malkova and Haber 2012; Lambert et al. 2005; Marie and Symington 2022).

The MIR2 pathway generates an insertion of ssDNA regions distant from the initiating DSB, between the two sites of invasion, as does MIR1 (**Fig. 2B**). Contrary to MIR1 however, MIR2 generates a single one-ended DSB used for product formation, precluding formation of secondary rearrangements. It uniquely requires displacement DNA synthesis and substantial sequence homology on the 3’ side of the donors, relative to the orientation of the invading strand (**Fig. 2B**). This sequence dependency may limit its broad applicability, likely confined to allelic recombination and large segmental duplications.

### Cellular state prone to non-conservative HR outcome

Coincident HR-mediated loss-of-heterozygosity (LOH) events have recently been reported (Sampaio et al. 2020), revealing the existence of an hyper-recombinogenic state characterized by crossover resolution in a subset of a yeast population. The nature and defect of this population remains unclear. Here, we show that resolution of MI JMs into a MIR1 translocation occurs late post-DSB induction, past the peak of D-loop and MI JMs formation (**Fig. 5G**). It may coincide with the onset of adaptation to the DNA damage checkpoint (Toczyski et al. 1997), at which Cdc5^PLK1^ phosphorylates a range of targets to trigger mitotic exit. These targets include Mus81-Mms4 (Matos and West 2014), an SSEs involved in crossover resolution (Ho et al. 2010), MIR (Piazza et al. 2017), and HR-dependent anaphase bridge cleavage in yeast and human cells (García-Luis and Machín 2014, Chan et al. 2018). Furthermore, aneuploidies of the initiating chromosomes are frequent (Piazza et al. 2017) (**Fig. 1E**), indicating that MIR is associated with chromosome mis-segregation. Consequently, cells undergoing several coincidental chromosomal rearrangements may be characterized by either (i) an inability to complete HR repair or dissolve DNA JMs prior to checkpoint adaptation, or (ii) an inefficient activation or rapid adaptation to the DNA damage checkpoint. Indeed, a crippled DNA damage checkpoint promotes LOH and SV formation in *C. glabrata*, which contributes to diversify the genome of this quasi-asexual yeast (Shor and Perlin 2021).

### Expanded sequence space for the generation of repeat-mediated SVs by HR

The prevailing model for the formation of HR-dependent balanced SVs between two repeated genomic elements is the canonical DSB repair (DSBR) model, which entails the endonucleolytic resolution of a dHJ formed between two ectopic repeats (Szostak et al. 1983). This model assumes a DSB will form within a repeat with sufficient homology on both sides to perform DNA strand invasion and second end capture (**Fig. 7A**). In *S. cerevisiae*, more than 300 bp on each side of the DSB is required for detectable crossover formation (Inbar et al. 2000). Conservation of this length dependence would rule out DSBR as a mechanism for *Alu*-mediated SV in humans. In contrast, MIR can be initiated by repeated sequences located away from the DSB site, provided that they are exposed by resection and part of the Rad51-ssDNA filament (**Fig. 7A**). Consequently, the sequence space at risk to form repeat-mediated SVs is expected to be much greater with MIR than with DSBR (**Fig. 7A**). Indeed, resection proceeds at 4 kb/hr (Zhu et al. 2008; Mimitou and Symington 2008) and can span tens of kilobases in mitotically dividing *S. cerevisiae*. Hence more than half of randomly distributed DSBs can undergo MIR after 2 hours of resection, despite a total repeat content of only 7% (Piazza and Heyer 2019). Likewise, resection reaches up to 3.5 kb in human U2OS cells (Zhou et al. 2014), which would make 70% of DSBs at risk of *Alu*-mediated SVs (**Fig. 7A**), the most frequently observed rearrangements in human tissues (Pascarella et al. 2022). This is ∼10-fold more than that expected from DSBR that requires the DSB to fall at a distance from the repeat edge (**Fig. 7A**). Comparatively shorter resection tracts in meiosis, averaging 0.8-1 kb both in *S. cerevisiae* and mouse (Mimitou et al. 2017; Zakharyevich et al. 2010; Yamada et al. 2020), may help protect against repeat-mediated SVs by MIR.

**Figure 7:**
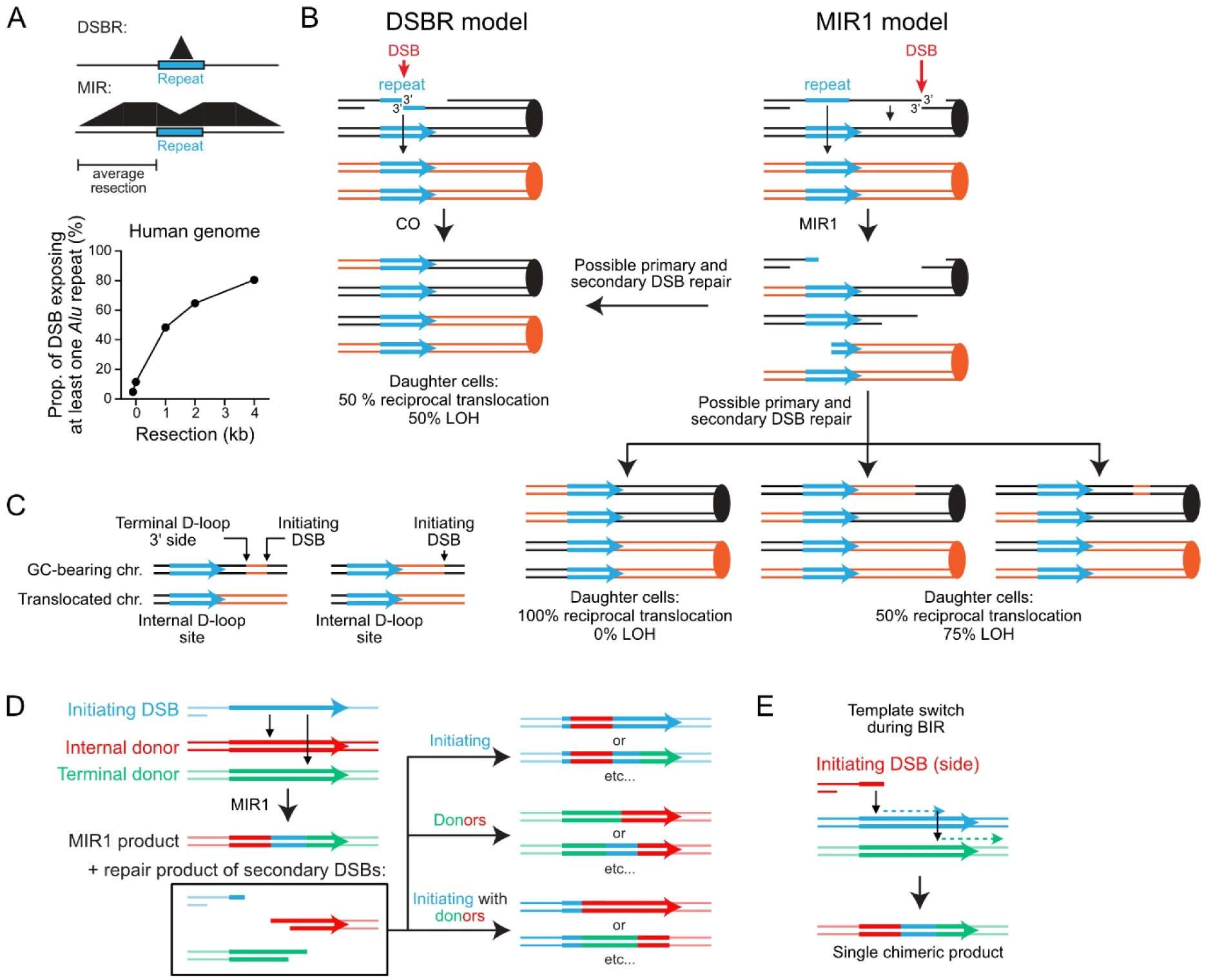
MIR and the formation of genome rearrangements. (A) Sequence space at risk of DSBR- and MIR-induced repeat-mediated SVs as a function of resection length. The example of *Alu* elements in the human genome is shown. (B) SV formation at an ectopic repeat according to the DSBR model (left) and MIR1 (right). DSBR requires the DSB and dHJs to form within the repeat, and in only half cases will segregation products cause LOH or a reciprocal SV. MIR1 requires the DSB to occur within the typical resection tract length from a repeat, and only requires one invasion at the repeat. Depending on the relative location and orientation of the two recombining repeats, MIR1 can produce circles, inversions, and tandem duplications and deletions. Final rearrangement and gene conversion outcomes depend on secondary DSB repair pathway, template used, and ends involved. (C) Repeat-mediated SV signature that can unambiguously be attributed to MIR. (D-E) MIR1 between repeats, and the repair of the initiating and secondary DSBs leads to several chimeric repeats. The co-occurrence of chimeric repeats can thus be used as a discriminating signature of MIR1, relative to template-switch during BIR that produces a single chimeric repeat.

Rearrangement patterns in yeast consistent with MIR have been reported. For instance, a break outside a Ty element is more prone to generate rearrangements than a DSB induced within (Hoang et al. 2010). Also, the recent observation that a large proportion of recombination events induced by DSBs within LTR or Ty elements have breakpoints in unique sequences and generate frequent recombination between non-targeted repeats (Qi et al. 2023; Fleiss et al. 2019) is consistent with the induction of secondary DSBs, as shown for MIR.

Of note and depending on the repair of the initiating and secondary DSB, MIR can elicit reciprocal translocations in 100% of daughter cells (**Fig. 7B**). Consequently, MIR1 is overall expected to lead to reciprocal translocation in >50% of cases, unlike DSBR.

### Repeat-mediated SV formation in human, and sequence signatures of MIR

Over the last decade, long-read sequencing technologies have allowed high confidence detection of balanced SVs in the human genome (Logsdon et al. 2020). A comparative analysis of multiple sequencing methodologies estimated that long-read sequencing detects seven-fold more variation in the form of insertions/deletions (indels) and SVs relative to high coverage Illumina whole-genome sequencing (Chaisson et al. 2019). Repeated elements, particularly *LINE* and *Alu* repeats, appears to be one of the primary sources of these *de novo* SVs, which are surprisingly common, even in healthy individuals (Chaisson et al. 2019; Balachandran et al. 2022; Pascarella et al. 2022). Yet much remains unexplored as to how repeat-mediated SVs contributes to human health and disease, and which specific HR sub-pathways are involved in their formation, namely DSBR, BIR, SSA, and MIR.

Here, we aim at providing sequence signatures that unambiguously distinguish SVs produced by MIR from that produced from other HR pathways (**Figs. 7C-E**). Indeed, repair of the secondary DSBs produced by MIR1 can lead to a reciprocal translocation at the repeat involved in MIR1 accompanied by a gene conversion of the intervening sequence between the repeat and the initiating DSB (**Fig. 7B**). In a subset of daughter cells, this repair outcome will lead to the association of a translocation on one chromosome and the presence of a flanking LOH on another chromosome (**Fig. 7B-C**). Of note is that both arise from the repair of secondary DSBs. The repeat-distal edge of the LOH tract reveals the position of the initiating DSB (**Fig. 7C**). If the LOH tract does not extend to the repeat, the repeat-proximal edge of the LOH indicates the 3’ side of the cleaved terminal D-loop (**Fig. 7C**). To our knowledge, no other mechanism than MIR1 can produce such matched SV/LOH tract at and flanking a repeated sequence.

Second, MIR1 can recombine long repeats such as *LINE* elements, inserting the sequence of the repeat flanking the initiating DSB between the translocated repeats (**Fig. 7D**). In a subset of cases, the initiating and secondary breaks will be repaired using the chimeric SV repeat, leading to the presence of two chimeric repeats (**Fig. 7D**). The presence of these matched chimeric repeats can distinguish MIR1 from a template-switch event during BIR, which is expected to produce only one chimeric repeat (**Fig. 7E**). Such signatures may help to refine mechanistic inferences made from long-read sequencing data and probe the associated genetic defects.

## Materials and methods

### Diploid *Saccharomyces cerevisiae* strains and genetic constructions

The genotype of the diploid *Saccharomyces cerevisiae* strains (W303 *RAD5^+^* background) used in this study are listed **Table S1**. Diploidy buffers for rearrangements that may cause loss of essential genes and allows for capture of more chromosomal rearrangements. Most genetic constructs have already been described in (Piazza et al. 2017). They are available as Genebank files in the **Dataset S1**. Briefly, strains contain a heterozygous copy of the *HO* endonuclease gene under the control of the *GAL1/10* promoter at the *TRP1* locus on chr. IV. The DSB-inducible construct contains the 117 bp HO cut-site (Fishman-Lobell and Haber 1992), various possible fragments of the *LYS2* gene lacking a start and a stop codons, and an unique 453 bp-long sequence derived from the PhiX genome. The DSB-inducible construct replaces the *URA3* locus (−16 to +855 from the start codon) on chr. V. The HO cut-site at the mating-type loci (*MAT*) on chr. III is inactivated by a point mutation to prevent HO cleavage (*MAT***a**-inc*/MAT*α-inc) (Fishman-Lobell and Haber 1992). In the allelic *inter-*chromosomal donor configuration, the *LY* and *S2* donors replace the original *LYS2* ORF on chr. II. In the ectopic-*cis* and -*trans* donor configuration, the *S2* donor and its 70 bp, 500 bp, or 1000 bp-long downstream sequence were inserted at the constitutively mutated *can1-100* locus, which caused a deletion of the beginning of the gene (−342 to +439 bp from the start codon). *S2* was oriented so as to avoid generating a dicentric II:V chromosome upon translocation with the *LY* donor left on chr. II.

The *RFC1-AID-9Myc::hphMX* construct is a gift from Neil Hunter and has been described previously (Kulkarni et al. 2020). The *POL3-iAID* construct is a gift from Lorraine Symington and has been described previously (Donnianni et al. 2019; Tanaka et al. 2015). The constructs for *TIR1* and *TetR* expression integrated at *SSN6* have been described previously (Tanaka et al. 2015).

### Media and culture conditions

Synthetic dropout (SD) and rich YPD (1% yeast extract, 2% peptone, 2% dextrose) solid and liquid media have been prepared according to standard protocols (Treco and Lundblad 2001). Liquid YEP-lactate (1% yeast extract, 2% peptone, 2% Lactate) were made using 60% Sodium DL-lactate syrup. All cultures were performed at 30°C.

### Pol3 and Rfc1 depletion using the auxin-inducible degron system

For time courses involving *pol3-iAID RFC1* or *pol3-iAID rfc1-AID-9Myc* strains, a single colony of the appropriate strain was used to inoculate a 5 mL YPD culture, grown to saturation overnight, and diluted in 250 mL YEP-lactate. Following ∼13-16 hours growth overnight at 30° C, the OD_600_ of the culture was determined, and the culture was split equally into two smaller flasks. Protein depletion was conducted as in (Donnianni et al. 2019) with minor modifications (**Fig. S7B**). 0.1 μg/mL doxycycline was added to the “with inhibitor” culture. One hour later, the culture was supplemented with 50 μg/mL doxycycline and 1.5 mM indole-3-acetic acid (IAA; 2 mM in (Donnianni et al. 2019)). An equal volume and concentration of solvent was added to the “without inhibitor” culture at both times (Donnianni et al. 2019). One hour later, the DSB was induced by addition of 2% galactose. Doxycycline and IAA were prepared as described in (Tanaka et al. 2015).

### Lysine prototrophy-based translocation assay in *S. cerevisiae*

The translocation assay has been described previously (Piazza et al. 2017, 2021b). Briefly, the basal Lys^+^ frequency was determined by plating yeast cells exponentially growing in YEP-lactate liquid culture on SD–LYS and YPD plates (control plating). The expression of the HO endonuclease was triggered in the remaining culture upon addition of 2% galactose. Two hours after galactose addition, when HO cutting is >99% (Piazza et al. 2021b, 2017), the induced Lys^+^ frequency is determined by plating again on SD–LYS and YPD plates. Basal and induced Lys^+^ frequencies as well as viability are determined after incubation of the plates at 30° C for 2-3 days. At least 3 independent replicates have been performed for each strain. Translocation frequencies are reported in **Table S2**.

### Southern blot analysis of the Lys^+^ recombinants

Independent Lys^+^ colonies were patched on SD-LYS plates, and their DNA was extracted from a saturating overnight 5 mL SD-LYS liquid culture. DNA was digested by *Hind*III (for the *inter*-chromosomal donor construct), *Pst*I (for the ectopic *S2*-70bp donor construct), or *Pst*I+*Eco*RI (for the ectopic *S2*-1000bp donor construct) 4 hours at 37° C and migrated overnight in 0.8% Agarose-LE (Affymetrix) in 1X TBE at 50 V. The DNA was transferred from the gel onto an Amersham Hybond-XL membrane (GE healthcare) following the manufacturer instructions (alkali protocol). The membrane was blocked with Church buffer (1% BSA, 0.25 M Na_2_HPO_4_ pH7.3, 7% SDS, 1 mM EDTA) for 2-3 hrs at 65° C. The *LY*, *S2*, or *LYS2* probes (2, 2, and 4 kb-long, respectively) together with Phage λ DNA (molecular ladder) were radio-labeled by random priming with P^32^-αdCTP (6,000 Ci/mmol; Perkin-Elmer) using the Decaprime II kit (Ambion Inc) and incubated with the membrane overnight at 65° C. After 3 – 5 washes for 10 min at 65° C (20 mM Na_2_HPO_4_ pH 7.3, 1% SDS, 1 mM EDTA), membranes were exposed for 8 to 24 hours, and the Storage Phosphor Screen (GE healthcare) scanned on a Storm Phosphorimager (Molecular Dynamics).

### Molecular karyotyping analyses

Preparation of DNA plugs and pulse-field gel electrophoresis (PFGE) of Lys^+^ recombinants was performed as described previously (Piazza et al. 2017). Array comparative genomic hybridization (aCGH) copy number profiling, and characterization of rearrangement junctions through Oxford Nanopore long-read sequencing were carried out following procedures described previously (Heasley et al. 2020; Zhang et al. 2013)

### D-loop Capture (DLC) and D-loop Extension (DLE) assays

The DLC and DLE assays have been described previously (Piazza et al. 2019, 2018) (for a step-by-step protocol, see (Reitz et al. 2022)). A psoralen crosslink reversal step adapted from (Yeung et al. 1988) has been added prior to the qPCR step, which removes various amplification biases and enables quantitative comparison of the amount of distinct JMs (Reitz et al. 2022). Briefly, psoralen crosslink reversal is performed upon incubation of purified DNA in decrosslinking solution (100 mM KOH, 10 mM Tris-HCl pH 8.0, 1 mM EDTA) at 90° C for 30 minutes. pH is neutralized by addition of 73 mM Na-Acetate and the DNA used for qPCR quantification. Primers used are listed in **Table S3**.

### Chromosomal Rearrangement-Capture (CR-C) assay

Genomic DNA was purified from 5 x 10^8^ cells collected at various time post-DSB induction by spheroplasting cells with Zymolyase, protein digestion, phenol-chloroform-isoamyl alcohol (25:24:1) extraction, isopropanol precipitation, and RNAse A treatment according to standard protocols. DNA was quantified on a Qubit 2.0 using the dsDNA HS Assay Kit according to manufacturer instructions. 500 ng of DNA was digested for 1 h at 37° C with 50 U of *Eco*RI-HF, and the enzyme was inactivated at 65° C for 20 min. 80 ng of digested DNA was ligated in 800 μL ligation buffer (50 mM Tris-HCl pH 8.0, 10 mM MgCl_2_, 10 mM DTT, 100 ng/μL BSA,1 mM ATP) with 20 U DNA ligase T4 for 1.5 h at 16° C. DNA was subsequently purified by phenol:chloroform:isoamyl alcohol (25:24:1) extraction followed by isopropanol precipitation. DNA was resuspended in 40 μL TE pH8 (10 mM Tris-HCl, 1 mM EDTA), and 2 μL used per qPCR reaction. It represents ∼10^5^ haploid genomes per reaction. Single-step qPCR amplification was performed on a Roche LightCycler 480 or a BioRad CFX96 thermocycler using the respective manufacturer’s SYBR green kit and instructions. The amplification of the rearranged molecule produced upon circularization of the 4,628 bp MIR1-containing DNA fragment was normalized on the average circularization efficiency of 3 DNA fragments of similar size (4,327, 4,701 and 4,888 bp). Primers used are listed in **Table S3**.

### Immunoblotting

Protein extracts for Western blot were prepared using a standard protocol for tricholoroacetic acid (TCA) extraction. ∼25 μL sample was run out on a 7.5% SDS-PA at 150V for ∼1 h alongside a protein standard (BioRad Precision Plus Protein Dual Color Standard). Samples were transferred to a PVDF membrane (BioRad TransBlot Turbo Mid-size LF PVDF membrane) using the BioRad Trans-Blot Turbo Transfer System. The membrane was then blotted using mouse anti-mini-AID-tag (1:1000) primary antibody (MBL), mouse anti-c-Myc (1:1000) primary antibody (SantaCruz Biotechnology), or mouse anti-GAPDH (1:10000) primary antibody (ThermoFisher), and anti-mouse HRP-conjugated (1:1000) secondary antibody (Agilent).

### Dilution series

Cell viability was assessed on YPD + solvent or YPD + 1.5 mM IAA, 50 μg/mL doxycycline as follows. 5 mL YPD cultures in 15 mL glass tubes were inoculated with a single colony corresponding to the appropriate strain and grown on a rotator at 30° C overnight. After ∼16 h of growth, the OD_600_ of the cultures was determined, and these cultures were used to inoculate fresh 5 mL YPD cultures at equivalent OD. The 5 mL cultures were grown on a rotator for ∼6 h at 30° C, then the OD_600_ of the cultures was determined again, and all cultures were diluted to an OD_600_ of 0.19. A series of 1:10 dilutions was prepared for each of the strains from these starting dilutions, and spotted onto YPD + solvent or YPD + 1.5 mM IAA, 50 μg/mL doxycycline plates in parallel using a multi-channel pipette. Plates were imaged following 2 days growth at 30° C.

### Calculation of the proportion of repeats at risk of SV formation by DSBR or MIR

The RepeatMasker BED track for the human genome assembly hg38 T2T CHM13v2.0 was obtained from the UCSC Table browser on 2022-10-19. Lines corresponding to *Alu* elements were retained, sorted with Bedtools sort, and overlapping elements merged with the Bedtools merge function, providing the proportion of the genome made of *Alu* elements (7%). Intervals were extended by 1, 2, and 4 kb, merged, and the genome fraction at risk of *Alu*-mediated SVs through MIR computed. Differently, initial intervals were shortened by 0.1 kb on each side to determine the genome fraction at risk of *Alu*-mediated SVs through DSBR, which requires the DSB to fall away from the repeat element edges.

## Acknowledgments

We are grateful to Lorraine Symington and Neil Hunter for providing Pol3 and Rfc1 depletion systems. D.R. was supported by T32CA108459 and a fellowship from the A.P. Giannini Foundation. Research in the Heyer laboratory is supported by grants GM58015 and GM137751. Research in the Piazza laboratory is supported by the European Research Council (ERC-StG 3D-loop, grant agreement 851006). Genome stability research in the Argueso laboratory is supported by NIH/NIGMS award R35GM11978801.

## Author contributions

Conceptualization: WDH and AP; Experiments: DR, YD, RAW, PR, and AP; Data analysis: DR, RAW, PR, and AP; Data interpretation: DR, RAW, JLA, WDH, and AP; Supervision: JLA, WDH, and AP; Funding Acquisition: JLA, WDH, and AP; Writing – original draft: AP; Writing – final draft: DR, WDH, and AP.

## References

1. Anand RP, Lovett ST, Haber JE. 2013. Break-induced DNA replication. Cold Spring Harb Perspect Biol 5: a010397.

2. Anand RP, Tsaponina O, Greenwell PW, Lee C-S, Du W, Petes TD, Haber JE. 2014. Chromosome rearrangements via template switching between diverged repeated sequences. Genes Dev 28: 2394–2406.

3. Argueso JL, Westmoreland J, Mieczkowski PA, Gawel M, Petes TD, Resnick MA. 2008. Double-strand breaks associated with repetitive DNA can reshape the genome. Proc Natl Acad Sci 105: 11845–11850.

4. Balachandran P, Walawalkar IA, Flores JI, Dayton JN, Audano PA, Beck CR. 2022. Transposable element-mediated rearrangements are prevalent in human genomes. Nat Commun 13: 7115.

5. Brocas C, Charbonnier J-B, Dhérin C, Gangloff S, Maloisel L. 2010. Stable interactions between DNA polymerase δ catalytic and structural subunits are essential for efficient DNA repair. DNA Repair 9: 1098–1111.

6. Burkovics P, Sebesta M, Sisakova A, Plault N, Szukacsov V, Robert T, Pinter L, Marini V, Kolesar P, Haracska L, et al. 2013. Srs2 mediates PCNA-SUMO-dependent inhibition of DNA repair synthesis. EMBO J 32: 742–755.

7. Chaisson MJP, Sanders AD, Zhao X, Malhotra A, Porubsky D, Rausch T, Gardner EJ, Rodriguez OL, Guo L, Collins RL, et al. 2019. Multi-platform discovery of haplotype-resolved structural variation in human genomes. Nat Commun 10: 1784.

8. Chan YW, Fugger K, West SC. 2018. Unresolved recombination intermediates lead to ultra-fine anaphase bridges, chromosome breaks and aberrations. Nat Cell Biol 20: 92–103.

9. Cosenza MR, Rodriguez-Martin B, Korbel JO. 2022. Structural variation in cancer: role, prevalence, and mechanisms. Annu Rev Genomics Hum Genet.

10. Deem A, Barker K, VanHulle K, Downing B, Vayl A, Malkova A. 2008. Defective break-induced replication leads to half-crossovers in *Saccharomyces cerevisiae*. Genetics 179: 1845– 1860.

11. Donnianni RA, Symington LS. 2013. Break-induced replication occurs by conservative DNA synthesis. Proc Natl Acad Sci U S A 110: 13475–13480.

12. Donnianni RA, Zhou Z-X, Lujan SA, Al-Zain A, Garcia V, Glancy E, Burkholder AB, Kunkel TA, Symington LS. 2019. DNA polymerase delta synthesizes both strands during break-induced replication. Mol Cell 76: 371–381.e4.

13. Fishman-Lobell J, Haber JE. 1992. Removal of nonhomologous DNA ends in double-strand break recombination: the role of the yeast ultraviolet repair gene *RAD1*. Science 258: 480–484.

14. Fleiss A, O’Donnell S, Fournier T, Lu W, Agier N, Delmas S, Schacherer J, Fischer G. 2019. Reshuffling yeast chromosomes with CRISPR/Cas9. PLOS Genet 15: e1008332.

15. Forget AL, Kowalczykowski SC. 2012. Single-molecule imaging of DNA pairing by RecA reveals a three-dimensional homology search. Nature 482: 423–427.

16. García-Luis J, Machín F. 2014. Mus81-Mms4 and Yen1 resolve a novel anaphase bridge formed by noncanonical Holliday junctions. Nat Commun 5: 5652.

17. Heasley LR, Watson RA, Argueso JL. 2020. Punctuated aneuploidization of the budding yeast genome. Genetics 216: 43–50.

18. Ho CK, Mazón G, Lam AF, Symington LS. 2010. Mus81 and Yen1 promote reciprocal exchange during mitotic recombination to maintain genome integrity in budding yeast. Mol Cell 40: 988–1000.

19. Hoang ML, Tan FJ, Lai DC, Celniker SE, Hoskins RA, Dunham MJ, Zheng Y, Koshland D. 2010. Competitive repair by naturally dispersed repetitive DNA during non-allelic homologous recombination. PLoS Genet 6: e1001228.

20. ICGC/TCGA Pan-cancer analysis of whole genomes consortium. 2020. Pan-cancer analysis of whole genomes. Nature 578: 82–93.

21. Inbar O, Liefshitz B, Bitan G, Kupiec M. 2000. The relationship between homology length and crossing over during the repair of a broken chromosome. J Biol Chem 275: 30833–30838.

22. Kulkarni DS, Owens SN, Honda M, Ito M, Yang Y, Corrigan MW, Chen L, Quan AL, Hunter N. 2020. PCNA activates the MutLγ endonuclease to promote meiotic crossing over. Nature 586: 623–627.

23. Lambert S, Watson A, Sheedy DM, Martin B, Carr AM. 2005. Gross chromosomal rearrangements and elevated recombination at an inducible site-specific replication fork barrier. Cell 121: 689–702.

24. Li X, Stith CM, Burgers PM, Heyer W-D. 2009. PCNA is required for initiation of recombination-associated DNA synthesis by DNA polymerase delta. Mol Cell 36: 704–713.

25. Liu J, Ede C, Wright WD, Gore SK, Jenkins SS, Freudenthal BD, Todd Washington M, Veaute X, Heyer W-D. 2017. Srs2 promotes synthesis-dependent strand annealing by disrupting DNA polymerase δ-extending D-loops. eLife 6: e22195.

26. Logsdon GA, Vollger MR, Eichler EE. 2020. Long-read human genome sequencing and its applications. Nat Rev Genet 21: 597–614.

27. Lydeard JR, Jain S, Yamaguchi M, Haber JE. 2007. Break-induced replication and telomerase-independent telomere maintenance require Pol32. Nature 448: 820–823.

28. Malkova A, Haber JE. 2012. Mutations arising during repair of chromosome breaks. Annu Rev Genet 46: 455–473.

29. Marie L, Symington LS. 2022. Mechanism for inverted-repeat recombination induced by a replication fork barrier. Nat Commun 13: 32.

30. Matos J, West SC. 2014. Holliday junction resolution: Regulation in space and time. DNA Repair 19: 176–181.

31. Mazón G, Lam AF, Ho CK, Kupiec M, Symington LS. 2012. The Rad1-Rad10 nuclease promotes chromosome translocations between dispersed repeats. Nat Struct Mol Biol 19: 964–971.

32. Mazón G, Symington LS. 2013. Mph1 and Mus81-Mms4 prevent aberrant processing of mitotic recombination intermediates. Mol Cell 52: 63–74.

33. McVey M, Khodaverdian VY, Meyer D, Cerqueira PG, Heyer W-D. 2016. Eukaryotic DNA polymerases in homologous recombination. Annu Rev Genet 50: 393–421.

34. Mimitou EP, Symington LS. 2008. Sae2, Exo1 and Sgs1 collaborate in DNA double-strand break processing. Nature 455: 770–774.

35. Mimitou EP, Yamada S, Keeney S. 2017. A global view of meiotic double-strand break end resection. Science 355: 40–45.

36. Miura T, Shibata T, Kusano K. 2013. Putative antirecombinase Srs2 DNA helicase promotes noncrossover homologous recombination avoiding loss of heterozygosity. Proc Natl Acad Sci 110: 16067–16072.

37. Notta F, Chan-Seng-Yue M, Lemire M, Li Y, Wilson GW, Connor AA, Denroche RE, Liang S-B, Brown AMK, Kim JC, et al. 2016. A renewed model of pancreatic cancer evolution based on genomic rearrangement patterns. Nature 538: 378–382.

38. Pardo B, Aguilera A. 2012. Complex chromosomal rearrangements mediated by break-induced replication involve structure-selective endonucleases. PLoS Genet 8: e1002979.

39. Pascarella G, Hon CC, Hashimoto K, Busch A, Luginbühl J, Parr C, Hin Yip W, Abe K, Kratz A, Bonetti A, et al. 2022. Recombination of repeat elements generates somatic complexity in human genomes. Cell 185: 3025–3040.e6.

40. Payen C, Koszul R, Dujon B, Fischer G. 2008. Segmental duplications arise from Pol32-dependent repair of broken forks through two alternative replication-based mechanisms. PLoS Genet 4: e1000175.

41. Piazza A, Bordelet H, Dumont A, Thierry A, Savocco J, Girard F, Koszul R. 2021a. Cohesin regulates homology search during recombinational DNA repair. Nat Cell Biol 23: 1176– 1186.

42. Piazza A, Heyer W-D. 2019. Moving forward one step back at a time: reversibility during homologous recombination. Curr Genet 65: 1333–1340.

43. Piazza A, Heyer W-D. 2018. Multi-invasion-induced rearrangements as a pathway for physiological and pathological recombination. BioEssays 40: 1700249.

44. Piazza A, Koszul R, Heyer W-D. 2018. A proximity ligation-based method for quantitative measurement of D-Loop extension in *S. cerevisiae*. Methods in Enzymology, Vol. 601, pp. 27–44.

45. Piazza A, Rajput P, Heyer W-D. 2021b. Physical and genetic assays for the study of DNA joint molecules metabolism and multi-invasion-induced rearrangements in *S. cerevisiae*. Methods in Molecular Biology, pp. 535–554.

46. Piazza A, Shah SS, Wright WD, Gore SK, Koszul R, Heyer W-D. 2019. Dynamic processing of displacement loops during recombinational DNA repair. Mol Cell 73: 1255–1266.e4.

47. Piazza A, Wright WD, Heyer W-D. 2017. Multi-invasions are recombination byproducts that induce chromosomal rearrangements. Cell 170: 760–773.e15.

48. Putnam CD, Kolodner RD. 2017. Pathways and mechanisms that prevent genome instability in *Saccharomyces cerevisiae*. Genetics 206: 1187–1225.

49. Qi L, Sui Y, Tang X-X, McGinty RJ, Liang X-Z, Dominska M, Zhang K, Mirkin SM, Zheng D-Q, Petes TD. 2023. Shuffling the yeast genome using CRISPR/Cas9-generated DSBs that target the transposable Ty1 elements. PLoS Genet 19: e1010590.

50. Reitz D, Savocco J, Piazza A, Heyer W-D. 2022. Detection of homologous recombination intermediates via proximity ligation and quantitative PCR in *Saccharomyces cerevisiae*. J Vis Exp e64240.

51. Robert T, Dervins D, Fabre F, Gangloff S. 2006. Mrc1 and Srs2 are major actors in the regulation of spontaneous crossover. EMBO J 25: 2837–2846.

52. Ruiz JF, Gómez-González B, Aguilera A. 2009. Chromosomal translocations caused by either pol32-dependent or pol32-independent triparental break-induced replication. Mol Cell Biol 29: 5441–5454.

53. Saini N, Ramakrishnan S, Elango R, Ayyar S, Zhang Y, Deem A, Ira G, Haber JE, Lobachev KS, Malkova A. 2013. Migrating bubble during break-induced replication drives conservative DNA synthesis. Nature 502: 389–392.

54. Sampaio NMV, Ajith VP, Watson RA, Heasley LR, Chakraborty P, Rodrigues-Prause A, Malc EP, Mieczkowski PA, Nishant KT, Argueso JL. 2020. Characterization of systemic genomic instability in budding yeast. Proc Natl Acad Sci U S A 117: 28221–28231.

55. Savocco J, Piazza A. 2021. Recombination-mediated genome rearrangements. Curr Opin Genet Dev 71: 63–71.

56. Shor E, Perlin DS. 2021. DNA damage response of major fungal pathogen *Candida glabrata* offers clues to explain its genetic diversity. Curr Genet.

57. Smith CE, Lam AF, Symington LS. 2009. Aberrant double-strand break repair resulting in half crossovers in mutants defective for Rad51 or the DNA polymerase delta complex. Mol Cell Biol 29: 1432–1441.

58. Stafa A, Donnianni RA, Timashev LA, Lam AF, Symington LS. 2014. Template switching during break-induced replication is promoted by the Mph1 helicase in *Saccharomyces cerevisiae*. Genetics 196: 1017–1028.

59. Szostak JW, Orr-Weaver TL, Rothstein RJ, Stahl FW. 1983. The double-strand-break repair model for recombination. Cell 33: 25–35.

60. Tanaka S, Miyazawa-Onami M, Iida T, Araki H. 2015. iAID: an improved auxin-inducible degron system for the construction of a ‘tight’ conditional mutant in the budding yeast *Saccharomyces cerevisiae*. Yeast 32: 567–581.

61. Toczyski DP, Galgoczy DJ, Hartwell LH. 1997. *CDC5* and *CKII* control adaptation to the yeast DNA damage checkpoint. Cell 90: 1097–1106.

62. Treco DA, Lundblad V. 2001. Preparation of yeast media. Curr Protoc Mol Biol Chapter 13: Unit13.1.

63. Wright WD, Heyer W-D. 2014. Rad54 functions as a heteroduplex DNA pump modulated by its DNA substrates and Rad51 during D loop formation. Mol Cell 53: 420–432.

64. Yamada S, Hinch AG, Kamido H, Zhang Y, Edelmann W, Keeney S. 2020. Molecular structures and mechanisms of DNA break processing in mouse meiosis. Genes Dev 34: 806–818.

65. Yeung AT, Dinehart WJ, Jones BK. 1988. Alkali reversal of psoralen cross-link for the targeted delivery of psoralen monoadduct lesion. Biochemistry 27: 6332–6338.

66. Zakharyevich K, Ma Y, Tang S, Hwang PY-H, Boiteux S, Hunter N. 2010. Temporally and biochemically distinct activities of Exo1 during meiosis: double-strand break resection and resolution of double Holliday junctions. Mol Cell 40: 1001–1015.

67. Zhang H, Zeidler AFB, Song W, Puccia CM, Malc E, Greenwell PW, Mieczkowski PA, Petes TD, Argueso JL. 2013. Gene copy-number variation in haploid and diploid strains of the yeast *Saccharomyces cerevisiae*. Genetics 193: 785–801.

68. Zhou Y, Caron P, Legube G, Paull TT. 2014. Quantitation of DNA double-strand break resection intermediates in human cells. Nucleic Acids Res 42: e19.

69. Zhu Z, Chung W-H, Shim EY, Lee SE, Ira G. 2008. Sgs1 helicase and two nucleases Dna2 and Exo1 resect DNA double strand break ends. Cell 134: 981–994.

