## Supplementary Figures and Tables for "Delineation of two multi-invasion-induced rearrangement pathways that differently affect genome stability"

### Supplementary information

#### Supplementary figure and table legends

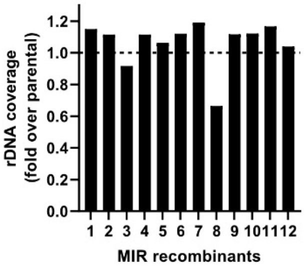

**Figure S1** (related to Figure 1): rDNA coverage was normalized onto the average genome-wide coverage and presented relative to the rDNA coverage in the parental strain (WDHY4260/APY89).

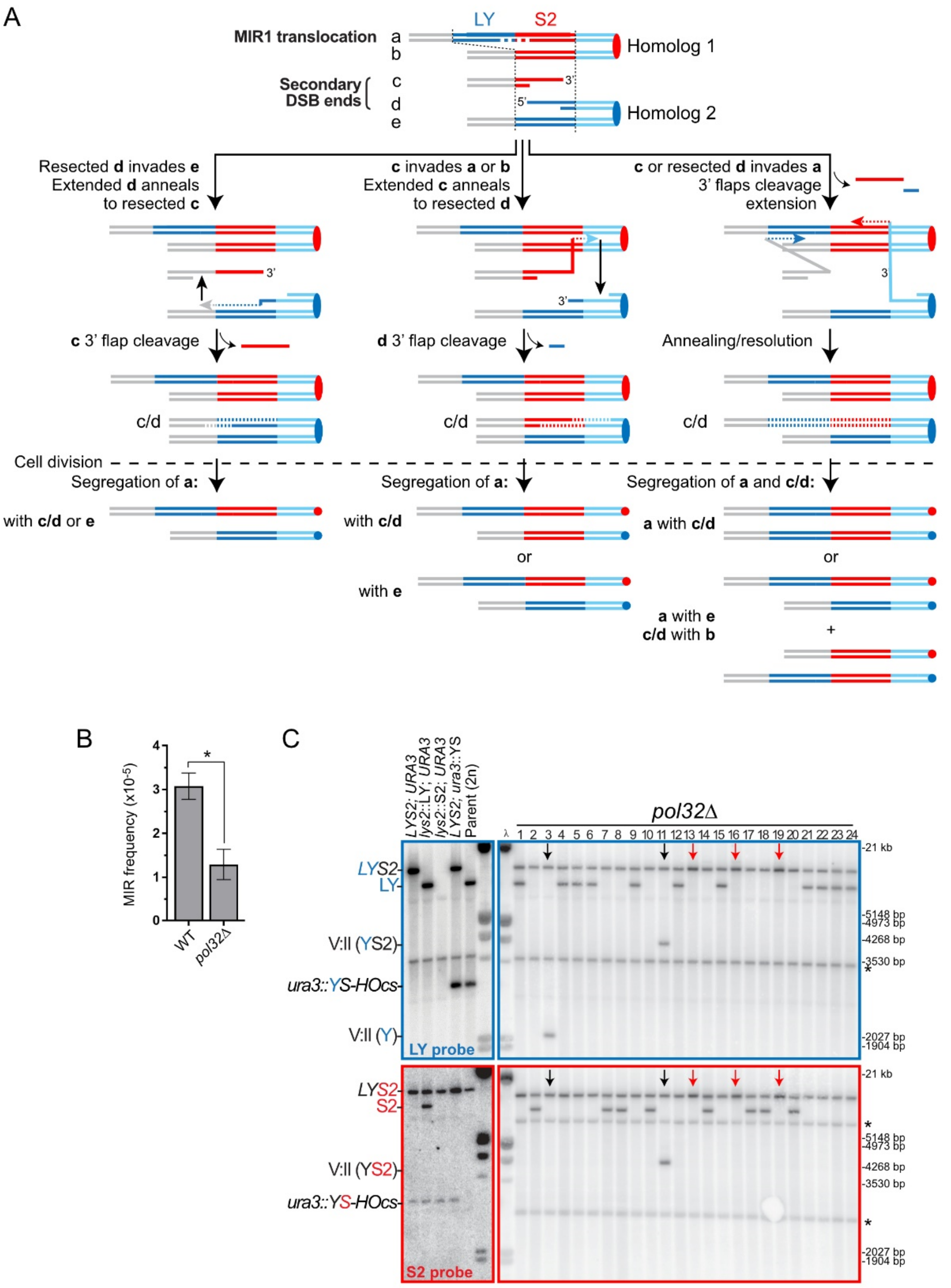

A
LY
S2
MIR1 translocation
a
b
Homolog 1
Secondary DSB ends
c
d
e
Homolog 2
Resected d invades e
Extended d anneals to resected c
c invades a or b
Extended c anneals to resected d
c or resected d invades a
3' flaps cleavage extension
c 3' flap cleavage
d 3' flap cleavage
Annealing/resolution
c/d
Cell division
Segregation of a:
Segregation of a:
Segregation of a and c/d:
with c/d or e
with c/d
or
with e
a with c/d
a with e
c/d with b
B
MIR frequency (x10-5)
WT
pol32Δ
C
LYS2; URA3
lys2::LY; URA3
lys2::S2; URA3
LYS2; ura3::YS
Parent (2n)
pol32Δ
LYS2
LY
V:II (YS2)
ura3::YS-HOcs
V:II (Y)
LY probe
S2
S2 probe
21 kb
5148 bp
4973 bp
4268 bp
3530 bp
2027 bp
1904 bp

Figure S2

**Figure S2** (related to Figure 2): (A) Model of secondary DSB repair following MIR and expected segregation outcomes. (B) MIR translocation frequency in wild-type and *pol32Δ* strains (WDHY4260 and WDHY4408, respectively; data from Piazza *et al.* 2017). (C) Southern blot analysis of 24 independent $Lys^+$ recombinants obtained in a *pol32Δ* mutant. Arrows indicate an anomaly in the segregation pattern, either because of additional rearrangements (black arrows) or presence of two *LYS2* translocation (red arrows; band intensity is two-fold higher than in other lanes). * indicates non-specific hybridization.

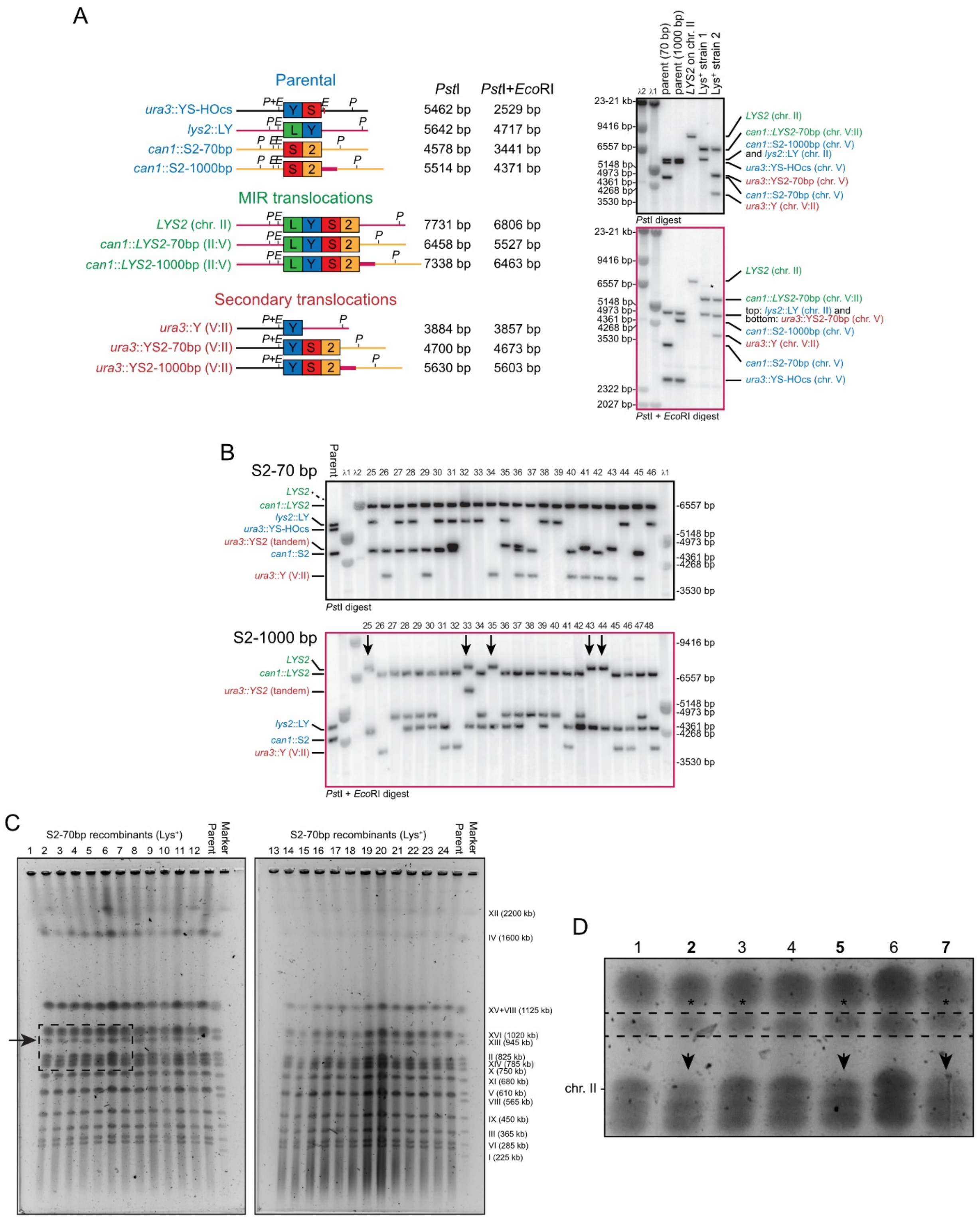
A
Parental
ura3::YS-HOcs
lys2::LY
can1::S2-70bp
can1::S2-1000bp
MIR translocations
LYS2 (chr. II)
can1::LYS2-70bp (II:V)
can1::LYS2-1000bp (II:V)
Secondary translocations
ura3::Y (V:II)
ura3::YS2-70bp (V:II)
ura3::YS2-1000bp (V:II)
PstI
PstI+EcoRI
5462 bp 2529 bp
5642 bp 4717 bp
4578 bp 3441 bp
5514 bp 4371 bp
7731 bp 6806 bp
6458 bp 5527 bp
7338 bp 6463 bp
3884 bp 3857 bp
4700 bp 4673 bp
5630 bp 5603 bp
parent (70 bp)
parent (1000 bp)
LYS2 on chr. II
Lys+ strain 1
Lys+ strain 2
PstI digest
PstI + EcoRI digest
LYS2 (chr. II)
can1::LYS2-70bp (chr. V:II)
can1::S2-1000bp (chr. V) and lys2::LY (chr. II)
ura3::YS-HOcs (chr. V)
ura3::YS2-70bp (chr. V)
can1::S2-70bp (chr. V)
ura3::Y (chr. V:II)
top: lys2::LY (chr. II) and bottom: ura3::YS2-70bp (chr. V)
can1::S2-1000bp (chr. V)
B
S2-70 bp
S2-1000 bp
Parent
LYS2
can1::LYS2
lys2::LY
ura3::YS-HOcs
ura3::YS2 (tandem)
can1::S2
ura3::Y (V:II)
C
S2-70bp recombinants (Lys+)
Parent
Marker
XII (2200 kb)
IV (1600 kb)
XV+VIII (1125 kb)
XVI (1020 kb)
XIII (945 kb)
II (825 kb)
XIV (785 kb)
X (750 kb)
XI (680 kb)
V (610 kb)
VIII (565 kb)
IX (450 kb)
III (365 kb)
VI (285 kb)
I (225 kb)
D
chr. II

Figure S3

**Figure S3** (related to Figure 2): (A) Structure, predicted length, and Southern blot validation of the DSB-inducible and donor substrates (blue), MIR translocation products (green) and secondary rearrangements products (red). (B) Southern blot analysis of recombinants #25-46 and #25-48 obtained with either limited (*S2*-70bp) or extensive (*S2*-1000bp) 3' flanking homologies, respectively. (C) PFGE analysis of recombinants #1-12 obtained with the *S2*-70bp and *S2*-1000bp donors, respectively. The inset shows subtle size difference for the neo-chromosome of ~950 kb, with higher migrating species associated with a decrease of chr. II signal (arrows).

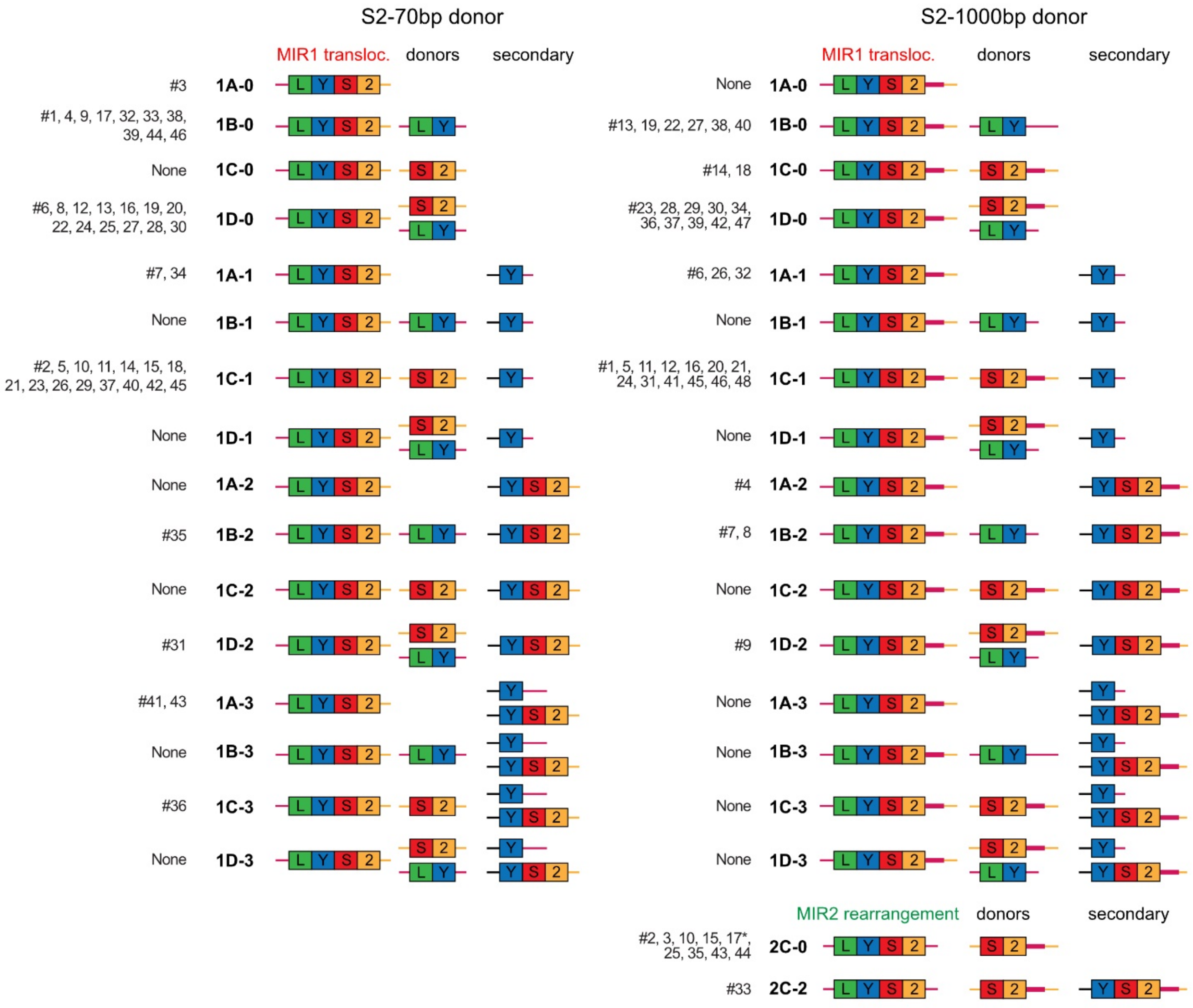

**Figure S4** (related to Figures 3 and 4): Structure of the MIR1 and MIR2 recombinants deduced from the Southern blot analysis with *S2*-70bp and *S2*-1000bp donors. * marks a composite strain bearing both a MIR1 and MIR2 product.

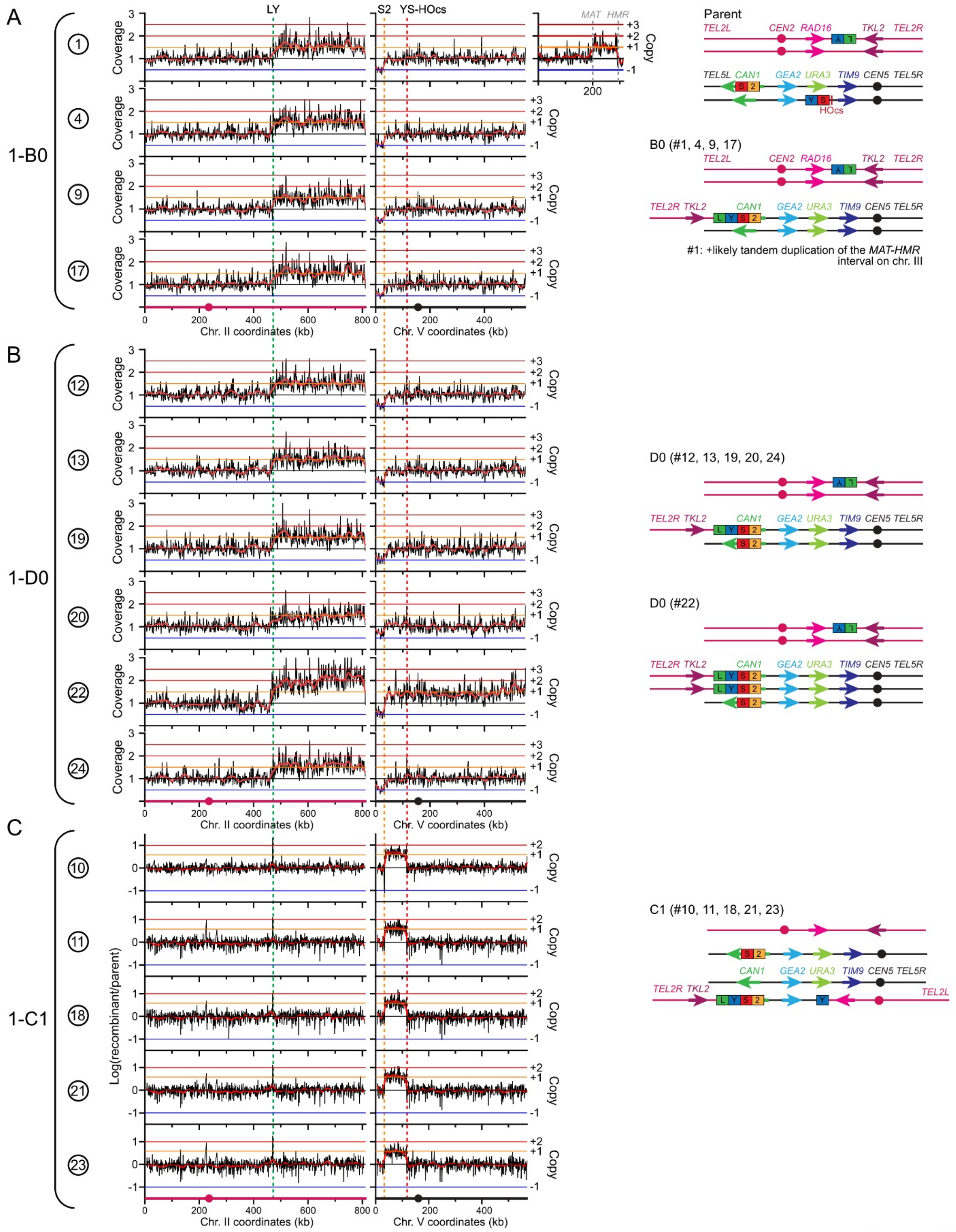

A
B
C
1-B0
1-D0
1-C1
LY
S2
YS-HOcs
MAT
HMR
Coverage
Copy
Log(recombinant/parent)
Chr. II coordinates (kb)
Chr. V coordinates (kb)
Parent
TEL2L CEN2 RAD16 TKL2 TEL2R
TEL5L CAN1 GEA2 URA3 TIM9 CEN5 TEL5R
HOcs
B0 (#1, 4, 9, 17)
TEL2R TKL2
#1: +likely tandem duplication of the MAT-HMR interval on chr. III
D0 (#12, 13, 19, 20, 24)
D0 (#22)
C1 (#10, 11, 18, 21, 23)

Figure S5

**Figure S5** (related to Figures 3 and 4): (A-C) Copy number analysis of four MIR1 recombinants obtained with the *S2*-70bp donor belonging to the B0 class (A), the D0 class (B), and the C1 class (C), and their deduced genomic structure. Recombinant #1 exhibited an additional unselected CNV spanning the *MAT-HMR* interval. Recombinant #22 exhibited two copies of the II:V translocated chromosome. Copy number profiles were obtained from paired-end sequencing data (A, B) or aCGH (C).

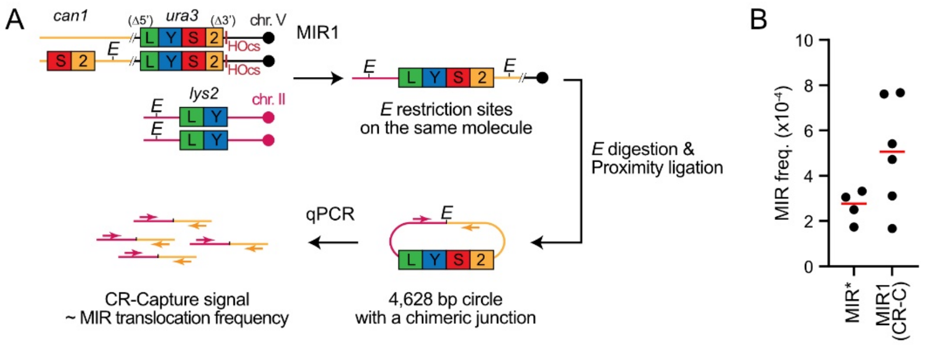

**Figure S6** (related to Figure 5): (A) Strain and procedure for MIR1 quantification by CR-C. The parental strain (APY625) contains a homozygous DSB-inducible construct at the *ura3* locus on chr. V, which purposefully precludes repair and growth resumption. It bears ~2 kb homology with ectopic donors: a homozygous *LY* donor at the *lys2* locus on chr. II and a heterozygous *S2* donor at *can1* on chr. V. A 152 bp unique sequence containing an *Eco*RI site was introduced 3' of the *S2*-70bp donor. (B) Comparison of induced MIR frequencies in wild-type cells determined genetically (*i.e.* $Lys^+$ colonies), and molecularly by CR-Capture 24 hours post-DSB induction. Parental strains differ in the number of DSB: MIR frequencies were obtained from a strain containing a heterozygous, repairable DSB site (APY611), while CR-Capture frequencies originated from a homozygous, unrepairable DSB site (APY625). * indicates that MIR values were divided by two in order to obtain a frequency per haploid genome equivalent.

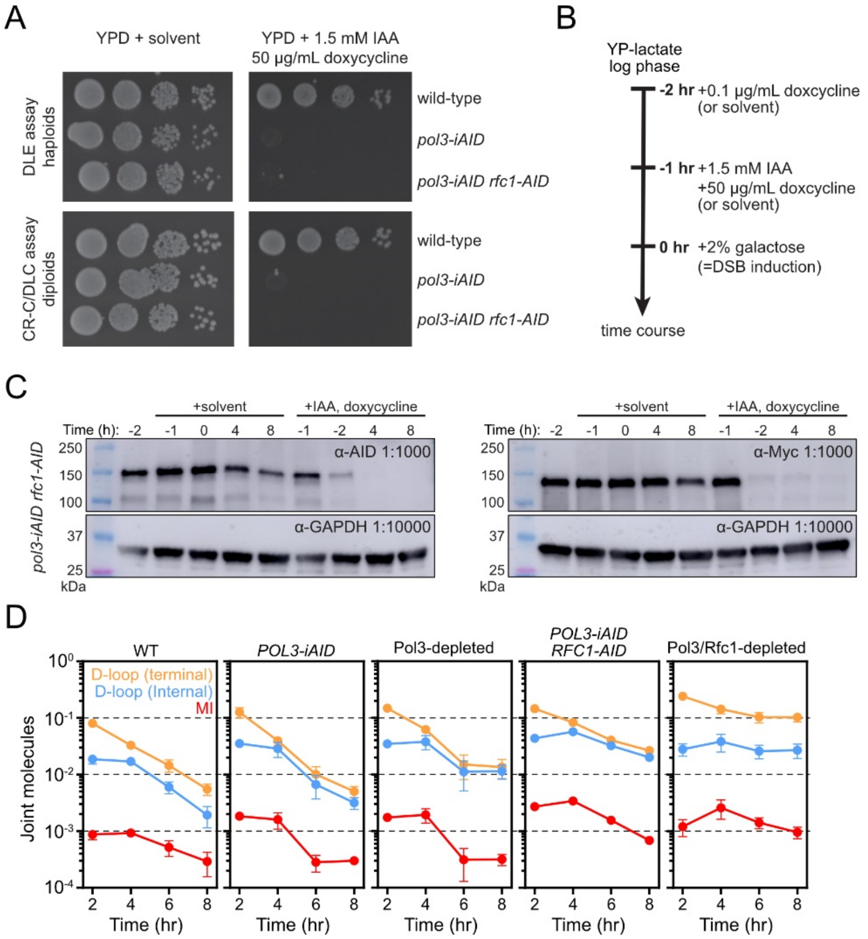

**Figure S7** (related to Figure 6): (A) Viability of wild-type, *pol3-iAID*, and *pol3-iAID rfc1-9Myc-AID* haploids used for DLE (APY266, WDHY6053, WDHY5067) and diploids used for CR-C (APY625, WDHY6065, WDHY6066) in rich media or inhibitor-containing media. Each spot corresponds to a 10-fold dilution. (B) Experimental scheme for Pol3 and Rfc1 depletion and DSB induction. (C) Western blot of untreated and inhibitor treated wild-type, *pol3-iAID*, and *pol3-iAID rfc1-9Myc-AID* haploids revealed with an anti-AID antibody (left; both proteins) and an anti-Myc antibody (right; Rfc1). Pol3-AID and Rfc1-9Myc-AID migrate at the same position. Time is indicated relative to DSB induction. (D) Absolute joint molecules detected by DLC in a wild-type strain (APY625, n≥8), in a *pol3-iAID* strain with and without inhibitors (WDHY6065, n=2), and in a *pol3-iAID rfc1-AID-9Myc* strain with and without inhibitor (WDHY6066, n=3).

### Table S1: Relevant genotype of *Saccharomyces cerevisiae* strains used in this study

| Table S1: Relevant genotype of *Saccharomyces cerevisiae* strains used in this study | | | |
|---|---|---|---|
| **Strain** | **Relevant genotype** | **Appears in Fig.** | **Source** |
| **Diploids** | | | |
| WDHY5144/APY89 | *MAT* **a**-*inc/MAT* α-*inc, ura3::YS(1000-1000)-HOcs/URA3, lys2::LY/lys2::S2, trp1::GAL-HO::hphMX/TRP1, his3 -11,15 /his3-11,15, can1-100/can1-100, ade2-1/ade2-1, leu2-3,112/leu2-3,112, RAD5/RAD5* | 1, 2C, S1 | This study |
| WDHY4408 | *WDHY5144, pol32::kanMX/pol32::kanMX* | 2C, S2 | Piazza *et al* . 2017 |
| WDHY5316/APY85 | *MAT* **a**-*inc/MAT* α-*inc, ura3::YS(1000-1000)-HOcs/ura3::loxP, can1::HIS3-S2-(70bp)-Pad/can1-100, lys2::LY/lys2::URA3, trp1::GAL-HO::hphMX/TRP1, his3 -11,15 /his3-11,15, ade2-1/ade2-1, leu2-3,112/leu2-3,112, RAD5/RAD5* | 3, 4, S2, S3, S4 | Piazza *et al* . 2017 |
| WDHY5317/APY86 | *WDHY5316, can1::HIS3-S2-500bp(trans)/can1-100* | 3A-B | This study |
| WDHY5318/APY87 | *WDHY5316, can1::HIS3-S2-1000bp(trans)/can1-100, lys2::LY/lys2::URA3* | 3, 4, S2A-B, S3 | This study |
| APY625 | *MAT* **a**-*inc/MAT* α-*inc, ura3::YS(2000-2000)-HOcs/ura3::YS(2000-2000)-HOcs, can1::HIS3-S2-Pad(EcoRI)/can1-100, lys2::LY/lys2::LY, trp1::GAL-HO::hphMX/TRP1, his3 -11,15 /his3-11,15, ade2-1/ade2-1, leu2-3,112/leu2-3,112, RAD5/RAD5* | 5A, 5E-G, 6A-B, S7D | This study |
| APY611 | *APY625, ura3::YS(2000-2000)-HOcs/ura3::loxP* | 5B, 5D | This study |
| APY704 | *APY625, rad51::kanMX/rad51::kanMX* | 5C | This study |
| WDHY6065/APY1354 | *APY625, pol3-iAID(kanMXN-term & hphMX C-term)/pol3-iAID(kanMX N-term & hphMX C-term), SSN6::pST1760 (TetR, OsTir1)::HIS3/SSN6::pST1760 (TetR, OsTir1)::HIS3* | 6A, S7A,C,D | This study |
| WDHY6066/APY1355 | *WDHY6065, RFC1-AID-9Myc::hphMX/RFC1-AID-9Myc::hphMX* | 6B, S7A,C,D | This study |
| **Haploids** | | | |
| APY266 | *MAT* **a**-*inc, ura3::LY-Hocs; lys2::LY; trp1::GAL-HO::hphMX, ade2-1, his3-11,15, leu2-3,112, can1-100, RAD5* | S7A | Piazza *et al* . 2021 |
| WDHY6053 | *APY266, pol3-iAID(kanMX N-term & hphMX C-term), SSN6::pST1760 (TetR, OsTir1)::HIS3* | 6A, S7A | This study |
| WDHY5067 | *WDHY6053, RFC1-AID-9Myc::hphMX* | 6B, S7A | This study |

### Table S2: Induced Lys$^+$ frequencies

| Table S2: Induced Lys$^+$ frequencies | | | | | | | | |
|---|---|---|---|---|---|---|---|---|
| **Genotype** | **DSB-inducible construct** | **Donors** | **Strain** | **Figure** | **Lys$^+$ frequency** | **SEM** | **n** | **Source** |
| WT | YS(1000-1000) | *inter* -chromosomal | WDHY5144/APY89 | **S2B** | 3.08E-05 | 3.00E-06 | 15 | Piazza *et al* . 2017 |
| *pol32 Δ* | YS(1000-1000) | *inter* -chromosomal | WDHY4408 | **S2B** | 1.29E-05 | 3.47E-06 | 4 | Piazza *et al* . 2017 |
| WT | YS(1000-1000) | Ectopic-trans; S2-70bp | WDHY5316/APY85 | **3B** | 2.80E-05 | 3.73E-06 | 6 | This study |
| WT | YS(1000-1000) | Ectopic-trans; S2-500bp | WDHY5317/APY86 | **3B** | 3.13E-05 | 4.90E-06 | 4 | This study |
| WT | YS(1000-1000) | Ectopic-trans; S2-1000bp | WDHY5318/APY87 | **3B** | 3.36E-05 | 5.26E-06 | 4 | This study |
| WT | YS(2000-2000) | Ectopic-trans; S2-70bp | APY611 | **S6*** | 5.30E-04 | 6.95E-05 | 4 | This study |
| * Represented divided by 2, to correct for ploidy for comparison with CR-C values | | | | | | | | |

### Table S3: Primers used in this study for the DLC, DLE, and CR-C assay.

#### DLC primers

| **Table S3: Primers used in this study.** | | |
|---|---|---|
| **Quantitative PCR primers** | | |
| **Name** | **Sequence (5'-3')** | **Purpose** |
| APO79/olWDH1760 | AGACAGAATTGGCAAAGATCC | Reference locus (*ARG4* , Ch.VIII); used as a dsDNA loading control. |
| APO80/olWDH1761 | GGCCAATTAGTTCACCAAGACG | |
| APO315/olWDH1762 | ACTTCGAATTTCGGCACTTC | Quantify the intra-molecular ligation efficiency of a 1904 bp *Eco* RI fragment at the *DAP2* locus. |
| APO316/olWDH1763 | CGATGAAACGTTAAGTGACCAC | |
| APO256/olWDH1766 | GTTTCAGCTTTCCGCAACAG | Measure dsDNA integrity at the HOcs. DSB induction causes a decrease of signal. |
| APO54/olWDH1767 | GGCGAGGTATTGGATAGTTCC | |
| APO596/olWDH2019 | CTTTAACCGGACGCTCGA | Quantify ssDNA at the PhiX site |
| APO597/olWDH2020 | TTGAGTTTATTGCTGCCGTC | |
| APO139/olWDH1764 | AGAGCGGTCAGTAGCAATCC | Quantify the DLC chimera formed between the upstream unique sequence of the invading molecule with the upstream unique sequence of the donor molecule at the intra-chromosomal (*CAN1* ) donor. |
| APO817 | GACGTACAAAGTTCCACTGGC | |
| APO139/olWDH1764 | AGAGCGGTCAGTAGCAATCC | Quantify the DLC chimera formed between the upstream unique sequence of the invading molecule with the upstream unique sequence of the donor molecule at the inter-chromosomal (*LYS2* ) donor. |
| APO319/olWDH1765 | CACACGCGAAAAACCGCC | |
| APO818 | AGTGTAGTTGGCCAAGTCATTC | Quantify the DLC chimera formed between the two donors in the MI intermediate. |
| APO319/olWDH1765 | CACACGCGAAAAACCGCC | |
| **Hybridization oligonucleotide** | | |
| **Name** | **Sequence (5'-3')** | **Purpose** |
| APO563/olWDH1770 | CGAAATCATCTTCGGTTAAATC CAAAACGGCAGAAGCCTGAATG AAACATATGAACCAATTGGAGG ACGTCAATGAATTCTGGGGATC CATTGCATTTTT | Hybridize at the *Eco* RI site upstream of the PhiX genome fragment on the resected broken molecule (chr.V). The last five nucleotides are not complementary. |

#### DLE primers

| Name | Sequence (5'-3') | Purpose |
|---|---|---|
| APO79/olWDH1760 | AGACAGAATTGGCAAAGATCC | Reference locus (*ARG4* , Chr. VIII); used as a dsDNA loading control. |
| APO80/olWDH1761 | GGCCAATTAGTTCACCAAGACG | |
| APO155/olWDH2052 | ATGTGCCTTCCTACCGCTC | Quantify the intra-molecular ligation (*i.e.* circularization) efficiency of a control 765 bp *Hin* dIII fragment (*YLR050C* ; Chr. XII). |
| APO156/olWDH2053 | TCAAGCGTGGTTACATTCCTTAC | |
| APO256/olWDH1766 | GTTTCAGCTTTCCGCAACAG | Measure dsDNA integrity at the HOcs. DSB induction causes a decrease of signal. |
| APO54/olWDH1767 | GGCGAGGTATTGGATAGTTCC | |
| APO596/olWDH2019 | CTTTAACCGGACGCTCGA | Quantify ssDNA at the PhiX site |
| APO597/olWDH2020 | TTGAGTTTATTGCTGCCGTC | |
| APO586/olWDH2009 | CACCACTTTGCCATTCAACAC | Quantify the chimera formed upon circularization of the extended invading molecule (DLE signal). To be normalized on the circularization efficiency. |
| APO587/olWDH2010 | TGCTCGGAGATTACCGAATC | |
| **Hybridization oligonucleotide** | | |
| **Name** | **Sequence (5'-3')** | **Purpose** |
| APO581/olWDH2007 | TCTGCTCGGAGATTACCGAATC AAAAAAATTTCAAAGAAACCGG AATCAAAAAAAAGAACAAAAA AAAAAAAGATGAATTGAAAAGC TTTATGGACCgac | Hybridize at the HindIII site upstream of the PhiX genome fragment on the resected broken molecule (Chr. V). The last three nucleotides are not complementary. |
| APO640/olWDH2046 | AATCTTTGTGAAGCTTCGCAAG TATTCATTTTAGACCCATGGTGG AACCCTAGTGTTGAATGGCAAA GTGGTGATAGAGTTCATAGAAT TGGTCAGTAT | Hybridize at the *Hin* dIII site downstream of the LY donor (Chr. II) on the extended invading molecule. |

#### CR-C primers

| Name | Sequence (5'-3') | Purpose |
|---|---|---|
| APO79/olWDH1760 | AGACAGAATTGGCAAAGATCC | Reference locus (*ARG4* , Chr. VIII); used as a dsDNA loading control. |
| APO80/olWDH1761 | GGCCAATTAGTTCACCAAGACG | |
| APO1331 | GATGAACTTCACCAGACAGC | Quantify the intra-molecular ligation (*i.e.* circularization) efficiency of a control 4,327p *EcoR* I fragment. |
| APO1332 | AACGTAGGGTCAAATTCAAGAG | |
| APO1325 | GCATGGATAAGCATCCTGAC | Quantify the intra-molecular ligation (*i.e.* circularization) efficiency of a control 4,701 bp *EcoR* I fragment. |
| APO1326 | GCTTGAATCTTGGCAGAAGAC | |
| APO1328 | CGCCTGGAATAAGCCAAAG | Quantify the intra-molecular ligation (*i.e.* circularization) efficiency of a control 4,888 bp *EcoR* I fragment. |
| APO1329 | CGCAAAACAACGACAAGAC | |
| APO1196 | CTTCTGCCTCGTGAACTTCC | Quantify the chimera formed upon circularization of the MIR1 translocation product (4,628 bp circle). To be normalized on the average circularization efficiency. |
| APO319/olWDH1765 | CACACGCGAAAAACCGCC | |

#### Supplementary Dataset S1

Annotated sequences of the genetic constructs used in this study in Genbank format.
